# ‘Loop tracing’ feedback reveals mechanisms that drive instabilities in resource-host-parasite dynamics

**DOI:** 10.64898/2026.03.17.712361

**Authors:** Eden J. Forbes, Spencer R. Hall

## Abstract

How and why do species interactions produce unstable dynamics? In the simplest models, the answers are straightforward. In the Rosenzweig-MacArthur predator-prey model, resource self-facilitation due to predation mortality triggers oscillations; in Lotka-Volterra competition, positive feedback from stronger interspecific competition underlies alternative states. However, when unstable dynamics arise with three or more species, ‘how’ and ‘why’ answers become more opaque. We propose that dissection of feedback loops, chains of direct species interactions, can answer these questions in meso-scale models. To demonstrate, we disentangle instabilities in epidemics using three variations of a general yet mechanistic resource-host-parasite model. Resources introduce destabilizing self-facilitation but also positive interspecific direct effects on propagule production and transmission rate. Those direct effects then produce instabilities through feedback loops. First, we ‘trace’ how resource self-facilitation catalyzes oscillations by weakening faster, shorter, lower levels of feedback relative to longer, slower feedback of the whole system. Then, we show how resource-dependent propagule yield introduces positive ‘cascade fueling’ feedback, creating an Allee threshold inhibiting invasion of parasites. In a third variant, we traced how both resource-dependent components produced those unstable dynamics and more complex behaviors, including a period-doubling route to chaos to which we apply a form of loop tracing. Hence, we show how and why direct, positive effects of resources modulate feedbacks underlying oscillations, Allee effects, and more during epidemics. We propose that loop tracing, a generally applicable method, could empower ecologists to glean much deeper insight into dynamics of species interactions.

## INTRODUCTION

What triggers a ‘simple’ ecological system of interacting species to oscillate? What causes an ecological system to develop alternative stables states? The answers to both questions involve feedback. Feedback captures the net effect of increased density of a species on its own growth rate. Negative feedback stabilizes; when species coexist at a stable equilibrium, that equilibrium exhibits negative feedback. In contrast, positive feedback at an equilibrium creates instability. Two simple, classic examples illustrate this point. First, in the Rosenzweig-MacArthur model of predator-prey dynamics (Rosenzweig & MacArthur, 1963; Rosenzweig, 1971), prey experience positive density-dependence from predators with a saturating functional response (‘crowing is good’ due to ‘safety in numbers’; Murdoch et al. 2003). When this predation component of prey fitness prevails over self-limitation, net positive feedback triggers predator-prey oscillations. In the second example, Lotka-Volterra competition can produce alternative stable states (Grover 1997). In this case, either one or the other competitor goes extinct; they cannot coexist because destabilizing interspecific competition exceeds stabilizing intraspecific competition. Hence, both classic models illustrate how different forms of positive feedback, produced via different biological mechanisms, cause populations to oscillate or exhibit alternative stable states. Both models produce highly transparent outcomes of species interactions via easily discerned mechanisms of feedback. In principle, then, the key to understanding instabilities hinges on finding the key feedback governing them.

However, a long-standing challenge remains: how can ecologists extend those insights into models of three or more species? One typical answer involves finding equilibria and calculating their stability numerically. Another approach is to simulate systems of species interactions and then observe and quantify their behavior. With both approaches, the same problem arises. While each can describe dynamical behaviors and the range of outcomes possible (the ‘what’), the mechanisms of ‘how’ and ‘why’ behaviors like oscillations or alternative states occur can remain obscured. At some level, those outcomes must involve feedback, too, but how and why? Here, we advocate for analysis of feedback loops (Levins 1975, Puccia and Levins 1985, Dambacher et al. 2003, Novak et al. 2016, Simons et al. 2022, Cortez 2024) to mechanistically dissect dynamical behaviors of models of intermediate complexity. Such models of a few interacting species or dimensions have enough realistic biology to generate unstable behaviors. To understand these behaviors, we ‘trace’ loops to find their genesis. By isolating chains of species interactions involved, ‘tracing’ allows us to pinpoint ‘how’ and ‘why’ an ecological system’s behavior changes.

To illustrate the promise of ‘loop tracing’, we consider both oscillations and alternative states produced by a family of resource-host-parasite models. These four-dimension (hereafter four ‘species’) models can produce unstable dynamics while remaining tractable enough to comprehensively trace feedback – a sweet spot of realism vs. transparency. The model family is inspired by a planktonic disease system of a host-grazer (*Daphnia dentifera*) which consumes propagules of an environmentally-transmitted parasite (the virulent fungus *Metschnikowia bicuspidata*) while foraging on primary producers (phytoplankton: Hall et al. 2007). Hence, foraging of the consumer is tied to parasite exposure, as it is in a variety of resource-host-parasite systems (e.g., nematodes consumed by ruminants; trematode eggs consumed by snails or amphibian hosts; viruses consumed by moth larvae). Non-linear (saturating) foraging behavior produces positive density-dependence of prey in predator-prey dynamics. Can non-linear foraging also explain why resource-host-parasite systems oscillate? How and why, and what role do parasites play? We also consider the possibility that production of infectious propagules from dead, infected hosts depends on resource availability, hence, consumption of resources. This biology applies to a variety of non-*Daphnia* systems as well (e.g., review in Cressler et al. 2014). Resource-dependent propagule production can produce Allee effects in other disease models (Smith et al. 2015; Borer et al. 2023). But, how and why should this positive, direct, interspecific link between resources and propagules produce positive feedback underlying alternative states?

To answer these questions, we test hypotheses linking positive direct effects of resources to instabilities in disease systems using loop tracing. We hypothesize that positive density-dependence of resources (hence self-facilitation) primarily causes oscillations during epidemics. However, this resource self-facilitation effect does not directly trigger oscillations, like in the simpler Rosenzweig-MacArthur model. Instead, because the disease system involves four ‘species’, epidemic oscillations arise when feedback strength in the whole system (wired through all four species, producing longer, slower feedback) exceeds the strength of feedback in loops through one, two, and three species (wired through shorter, faster loops; Puccia and Levins 1985; Lever et al. 2022). Then, we hypothesized that the positive, direct, interspecific links from resources to propagules produce alternative stable states. By tracing out feedback created by this resource-propagule link, we isolated the key chains of species interactions that cause ‘cascade fueling’ that ultimately create Allee effects. In both cases, loop tracing provided mechanistic insight into dynamical behaviors that could otherwise be described, but not easily explained and understood. Finally, we demonstrate a nascent method for extending loop analysis to the stability of orbits at a period doubling event that occurs when we combine both mechanisms of instability. We hope that future loop tracing will uncover deeper mechanistic insights into behaviors of new models of intermediate complexity and tractability.

## MODEL VARIANTS AND THEIR FOUR LEVELS OF FEEDBACK

### Three model variants of resource-host-parasite interactions (Table 1)

Three model variants combine two ways in which resources can introduce potentially destabilizing positive feedback into host-parasite dynamics, first via resource clearance by grazer-hosts and second via resource-dependent propagule yield. All variants represent a freshwater system in which a zooplankton grazer-host (*Daphnia dentifera*) consumes algal resources distributed in the water column. Hosts become infected after inadvertently consuming spores of the virulent fungal parasite (*Metschnikowia bicuspidata*) while grazing. Based on this biology, each variant tracks dynamics of four species/classes (hereafter, ‘species’; see equations in Table 1, Fig. A1A). The general model structure is:

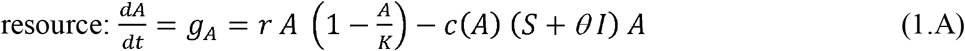

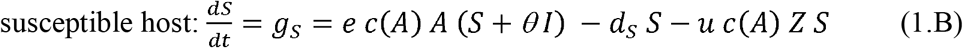

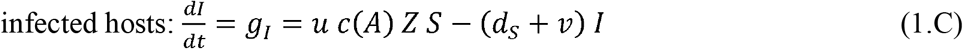

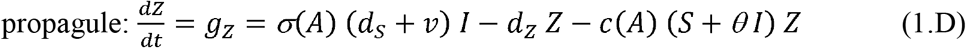

where *g*_*j*_ is the population growth rate of species *j*. The autotroph resource (*A*) grows logistically, with maximum per capita growth rate *r*, and carrying capacity *K* (eq. 1.A). The grazer-hosts consume that resource as governed by clearance rate *c*(*A*). ‘Clearance rate’ as described in predator-prey theory represents the per prey (resource, spore) risk of being eaten; it is also the amount of habitat ‘cleared’ of prey per unit time. Clearance rate is reduced virulently by infection when *θ* < 1 (Hite et al. 2017; Strauss et al. 2019; Penczykowski et al. 2022). Births of susceptible hosts (*S*) from both host classes (i.e., horizontal transmission) follow conversion of consumed resources, *c*(*A*) *A*, with efficiency *e* (eq. 1.B). Susceptible hosts die at background rate *d*_*S*_ and move to the infected class as they contact propagules (*Z*). Because hosts consume propagules, exposure is proportional to clearance rate *c*(*A*) (Hall et al. 2007), and infection ensues with per propagule infectivity *u*. Infected hosts (*I*) die at virulence-enhanced rate *d*_*S*_ *+ v*, then release *σ*(*A*) propagules into the environment (eq. 1.C). Hosts do not recover from infection. Free-floating propagules are removed at background rate *d*_*Z*_ or become consumed by both classes clearance rate (eq. 1.D).

**Table 1:** Summary of family of model variants with resources, hosts, and parasite (*ASIZ*; eq. 1), explanations of variables and parameters, with default values (see also Fig. A1A).

| *Term* | *Units* | *Definition* | *Value/Range* |
| --- | --- | --- | --- |
| **State variables** |  |  |  |
| *S* | hosts·L<sup>-1</sup> | Susceptible host density | - |
| *I* | hosts·L<sup>-1</sup> | Infected host density | - |
| *Z* | spores (sp)·L<sup>-1</sup> | Propagule density | - |
| *A* | mg·L<sup>-1</sup> | Algal resource density | - |
| **Parameters** |  |  |  |
| *c(A)* | L·host<sup>-1</sup>·day<sup>-1</sup> | Clearance rate function, aka,<br>exposure rate, per prey risk of<br>being eaten | Variable |
| *d<sub>S</sub>* | day<sup>-1</sup> | Background death rate, hosts | 0 – 0.15 |
| *d<sub>Z</sub>* | day<sup>-1</sup> | Loss rate, propagules | 0.9 |
| *e* | host·mg<sup>-1</sup> | Conversion efficiency | 8 |
| *f* | L·host<sup>-1</sup>·day<sup>-1 a</sup><br>or mg·host<sup>-1</sup>·day<sup>-1 a</sup> | Feeding rate – fixed slope or<br>maximum | 0.0153 |
| *h* | mg·L<sup>-1</sup> | Half-saturation constant, feeding<br>rate | 0.2314 |
| *K* | mg·L<sup>-1</sup> | Carrying capacity, resource | 0 – 6 |
| *o* | sp·host<sup>-1 b</sup><br>sp·day·mg<sup>-1 b</sup> | Propagule yield (x 10<sup>2</sup>) – fixed<br>or an efficiency<sup>b</sup> | 483.8 |
| $r$ | $\text{mg}\cdot\text{day}^{-1}$ | Max per capita growth rate,<br>resource | 1 |
| $u$ | $\text{host}\cdot\text{sp}^{-1}$ | Per propagule infectivity ($\times 10^{-2}$) | 1.049 |
| $v$ | $\text{day}^{-1}$ | Virulent mortality | 0.05157 |
| $\sigma(A)$ | $\text{sp}\cdot\text{host}^{-1}$ | Propagule yield (function) | Variable |
| $\theta$ | -- | Reduction of feeding rate,<br>infected hosts | 0.8 |

<sup>a</sup> Units differ for type I (top) vs. type III (bottom) clearance.
<sup>b</sup> Units and meaning differ: for fixed yield (variant 1), $\sigma(A) = o$ , where $o$ is a maximum yield (with units $\text{sp}\cdot\text{host}^{-1}$ ).
For linear clearance, where $c(A) = f$ , $\sigma(A) = o f A$ , so $o$ is an efficiency with units $\text{sp}\cdot\text{day}\cdot\text{mg}^{-1}$ , and $\sigma(A)$ increases
with $A$ . For type III clearance, $c(A) = f A / (h^2 + A^2)$ , so $\sigma(A) = o f A^2 / (h^2 + A^2)$ , so $o$ is again an efficiency with units
$\text{sp}\cdot\text{day}\cdot\text{mg}^{-1}$ . Now, $o f$ is a maximum yield ( $\text{sp}\cdot\text{host}^{-1}$ )

We made the three variants of this basic model structure by changing resource-dependence of clearance rate, *c*(*A*), and propagule yield, σ(*A*). Both these changes introduced or altered direct effects between species that could promote positive feedback. These direct effects, grouped in each model’s Jacobian matrix (Appendix section 1), denote how a species’ growth responds to changes in each species density in the model (e.g., direct effect ‘*J*_*XY*_’, denotes the consequence of an increase in species *Y* on the growth of species *X*). Notably, direct effects do not change stability alone, but rather in combination with one another (see below).

First, we considered the difference between linear vs. type III clearance:

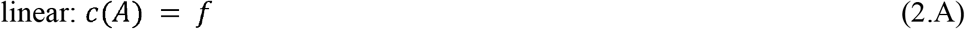

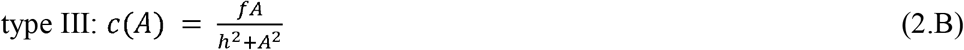

With linear clearance, the per resource or per spore risk of being eaten depends on constant rate *f*. In the type III functional response (Uszko et al. 2015), that risk is now unimodal with resource density, *A*, as governed by *f*, now the maximum feeding rate, and half-saturation constant, *h*. Type III clearance alters self-regulation of the resource, or the direct effect of the resource on itself (*J*_*AA*_). Type III clearance potentially introduces positive density-dependence at the individual level, where density-dependence is the sensitivity (slope) of fitness, aka per capita growth rate (*r*_*A*_ = *G*_*A*_/*A*), with an increase in the resource (so *DD*_*A*_ = ∂*r*_*A*_/∂*A*). Self-regulation is the sensitivity of intraspecific population growth rate to density (so, *J*_*AA*_ = ∂*g*_*A*_/∂*A = DD*_*A*_ *A +r*_*A*_); *J*_*AA*_ > 0 implies self-facilitation, while *J*_*AA*_ < 0 means self-inhibition. At an interior equilibrium, fitness of the resource is zero (therefore: *J*_*AA*_ *= DD*_*A*_ *A*), so a resource experiencing positive density dependence (*DD*_*A*_ > 0) thereby self-facilitates (*J*_*AA*_ > 0). With these definitions,

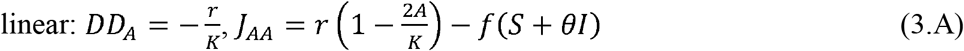

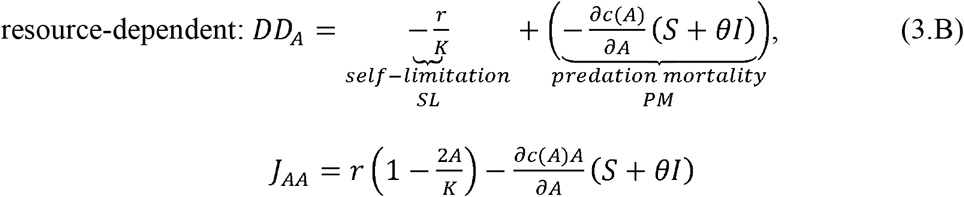

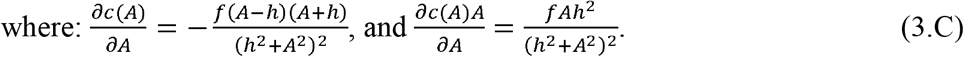

When the resource consumed by a host with type III clearance rate experiences positive density-dependence, the self-limitation component (*DD*_*SL*_ < 0) created by logistic growth (costs of crowding) becomes overmatched by the benefits of crowding from the ‘safety in numbers’ (*DD*_*PM*_ > 0, which requires *A* > *h*; eq. 3.A,C). In this case, the resource self-facilitates (*J*_*AA*_ > 0). Type III clearance in variant 1 also introduced a direct, interspecific effect from resources to infected host (*J*_*IA*_) because exposure is proportional to clearance rate.:

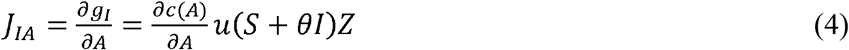

which is positive if *A* < *h*.

Second, we considered constant vs. resource-dependent propagule yield: constant:

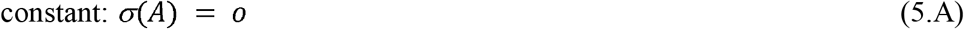

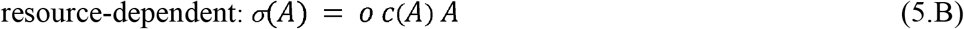

where in variant 1, *o* is a constant number of propagules released upon death of infected hosts (eq. 5.A). With linear clearance, yield increases with *A* with slope *o f*; with type III clearance, the maximum yield is *o f*. Conversely, resource-dependent yield in variants 2 and 3 increases proportionally with resource consumption, *c*(*A*) *A*, with *o* scaling the yield (as a conversion factor; eq. 5.B). In both variants, resource-dependent propagule yield introduced a new, potentially positive, interspecific direct effect of resources on propagules (*J*_*ZA*_). Since variant 2 assumed a linear clearance (with propagule yield of *o f A*) while variant 3 assumed type III clearance (with propagule yield of *o f A*^2^ / [*h*^2^ + *A*^2^]), we have, for:

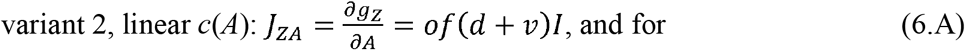

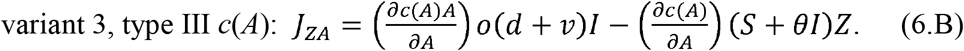

Resources increase growth rate of propagules (*J*_*ZA*_ > 0 always in eq. 6.A, and almost always in eq. 6.B). As we will show below, *J*_*ZA*_ introduces new feedback loops, including positive ‘cascade fueling’ loops, that can ultimately cause alternative states with Allee effects). Since the third variant combines type III clearance with resource-dependent propagule yield, it also allows the two forms of positive feedback to manifest and interact via *J*_*AA*_, *J*_*IA*_, and *J*_*ZA*_. Hence, as will be shown, variant 3 can yield both oscillations and simple alternative stable states with Allee effects, but also complex dynamics such as two new regions of bistability, period-doubled oscillations, and chaos.

### Feedback loops in the three model variants (and the genesis of instabilities in general; Fig. 1)

Positive intra (*J*_*AA*_) and interspecific (*J*_*IA*_, *J*_*ZA*_) direct effects of resources can alter system stability via feedback loops. Feedback loops describe how a small increase in density of a species would increase or decrease its growth rate via positive or negative feedback, respectively. Feedback loops are constructed by connecting species via chains of direct effects. Since our family of models has four equations, each variant generates four levels of feedback (Fig. 1A). Those familiar with Routh-Hurwitz criteria for stability will recognize these levels as the negative of coefficients of the characteristic polynomial of a system’s Jacobian matrix. The first level of feedback, *F*_1_, sums the intraspecific, self-regulatory, direct effect of each species on itself (Fig. 1B). Susceptible (*J*_*SS*_) and infected hosts (*J*_*II*_) and propagules (*J*_*ZZ*_) each produce self-regulation (i.e., in increase in their density would depress their growth rate, *J*_*SS*_ = ∂*G*_*S*_/∂*S* < 0, etc.). However, the resource, as explained above, can produce self-facilitation (*J*_*AA*_ > 0). In a two-dimensional model (like the Rosenzweig-MacArthur predator-prey model), *F*_1_ > 0 alone would trigger oscillations. However, with three or more ‘species’ (dimensions), oscillations start when the ‘Hopf criteria’ (*HC*) becomes positive; assuming each level of feedback is negative, for the four species models here, *HC* is:

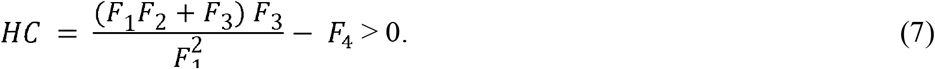

*HC* pits feedback at levels one (*F*_1_), two (*F*_2_), and three (*F*_3_) against the highest level (*F*_4_). Thus, self-facilitation of the resources (*J*_*AA*_ > 0) would trigger oscillations via its roles in influencing the magnitude of lower levels of feedback vs. the top-most in *HC*.

**Figure 1.**
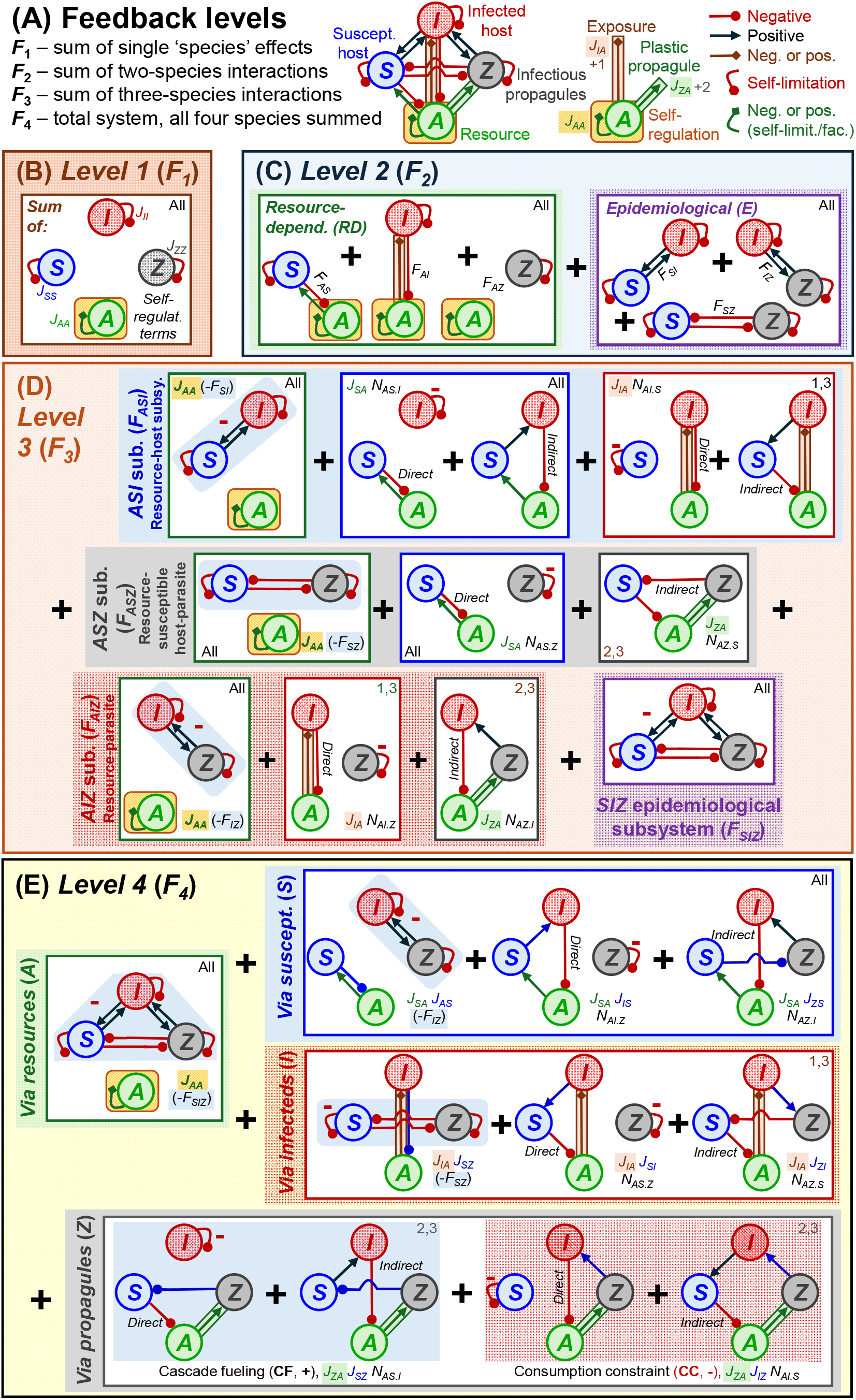
Four levels of feedback link resources to instabilities in resource-host-parasite dynamics. **(A)** A family of three model variants link resources (*A*), susceptible (*S*) and infected (*I*) hosts, and propagules (*Z*) through negative (circle), positive (arrow), or variably signed (diamond) direct effects. Resources introduce potentially positive direct effects through self-regulation (from *A* on itself, *J*_*AA*_, variants 1 & 3), through exposure via resource-dependent clearance (from *A* to *I, J*_*IA*_, 1 & 3), and through propagule release from infected hosts (from *A* to *Z, J*_*ZA*_, 2 & 3). Those direct links then contribute to one, two, three, and four-’species’ loops. **(B)** *Level 1 feedback* (*F*_1_) sums self-inhibition or facilitation effects of each of the four species. **(C)** *Level 2 feedback* (*F*_2_) sums binary interactions of three ‘resource-dependent’ pairs (each including *J*_*AA*_) and three epidemiological pairs involving hosts and propagules. **(D)** *Level three feedback* (*F*_3_) sums four sets of three-species interactions, three of which involve resources (i.e., feedback in *ASI, ASZ*, and *AIZ* subsystems) and one of which involves epidemiology only (*SIZ*). Finally, **(E)** *level four feedback* (*F*_4_) sums all four-species interactions. Starting with resources, the loops operate ‘via’ resources, susceptible hosts, infected hosts, and propagules. As shown, the four levels of feedback reflect the full model (variant 3) with all resource effects (*J*_*AA*_, *J*_*IA*_, *J*_*ZA*_). Model variants would sever some links (i.e., variant 1 disconnects *J*_*ZA*_, variant 2 severs *J*_*IA*_), thereby eliminating some loops (*via propagules* and *via infected hosts*, respectively). ‘Base’ loops sum the *via resource* and *via susceptible host* components.

Consequently, we propose to use ‘loop tracing’ to pinpoint if and how self-regulation of the resource (*J*_*AA*_), a direct intraspecific effect, creates oscillations by weakening chains of interactions involved in lower levels of feedback (*F*_1_, *F*_2_, *F*_3_) relative to *F*_4_. In this way, *J*_*AA*_ mirrors but modifies its role in classic predator-prey models. *J*_*AA*_ is found throughout levels of feedback. Level two feedback (*F*_2_) sums feedback in binary pairs (Fig. 1C). Three of those pairs are ‘resource-dependent’ (*RD*) – they contain the resource, hence *J*_*AA*_, in ‘subsystems’ involving susceptible hosts (*F*_*AS*_), infected hosts (*F*_*AI*_), and propagules (*F*_*AZ*_). For example, feedback in the *F*_*AS*_ system is the difference of the product of interspecific effects (*J*_*AS*_ *J*_*SA*_) and intraspecific ones (*J*_*AA*_ *J*_*SS*_; so *F*_*AS*_ = *J*_*AS*_ *J*_*SA*_ – *J*_*AA*_ *J*_*SS*_; see Appendix section 3 for more details). The other three ‘epidemiological’ (*E*) pairs relate susceptible host as the epidemiological resource to the two parasite life stages (in subsystems *F*_*SI*_, *F*_*SZ*_, and *F*_*IZ*_). If *J*_*AA*_ weakens level two feedback to help trigger oscillations, it will work via some set of the three resource-dependent subsystems (Fig 1C).

Level three feedback (*F*_3_) sums feedback in four trios of interacting species. Resources feature in the first three: the resource (*A*) - susceptible (*S*) – infected (*I*) subsystem (*F*_*ASI*_, blue box), then resources with propagules (*Z*) and one of the two host classes (*F*_*ASZ*_ [grey] and *F*_*AIZ*_ [red]; Fig. 1D). Each follows a similar structure, as illustrated by the resource-host subsystem (*F*_*ASI*_). Here, feedback among these three players starts, as derived from a ‘resource-centered’ perspective, with the effect of the resource on itself (*J*_*AA*_) times feedback in the *SI* subsystem (-*F*_*SI*_ [as described above]). The second part of *F*_*ASI*_ starts with the effect of the resource on *S* (*J*_*SA*_), then multiplies that by the net effect of *S* back onto *A* via *I, N*_*AS*.*I*_ (completing the three-species feedback). This three species net effect, in turn, sums direct and indirect components; the direct component is the effect of *S* back onto *A* directly, multiplied by self-regulation of *I* (*J*_*AS*_ [-*J*_*II*_]), while the indirect chain moves from *S* to *I* then back to *A* (*J*_*IS*_ *J*_*AI*_; so, *N*_*AS*.*I*_ = *J*_*AS*_ [-*J*_*II*_] + *J*_*IS*_ *J*_*AI*_). Analogous to the second part (*J*_*SA*_ *N*_*AS*.*I*_), the third part of *F*_*ASI*_ starts with the effect of *A* on *I* (*J*_*IA*_), then adds the net effect of *I* back on to *A* via *S* (*N*_*AI*.*S*_). Note that this third part requires non-zero interspecific effect of *A* on *I* (introduced via type III-based exposure rate in variants 1 and 3: *J*_*IA*_ ≠ 0); for variant 2 with linear clearance rate, the resource has no direct effect on infected hosts (*J*_*IA*_ = 0), wiping out this part of *ASI* subsystem feedback. The other two subsystems with resources (*F*_*ASZ*_, *F*_*AIZ*_) are composed similarly to *F*_*ASI*_ (Fig. 1D, Appendix Section 3). The fourth trio, the epidemiological subsystem (*F*_*SIZ*_), has no relationship to resources (Appendix section 3). Hence, we hypothesize that, if self-regulation of the resource (*J*_*AA*_) weakens level three feedback (*F*_3_), it will work via one or more of the trios with resources (*F*_*ASI*_, *F*_*ASZ*_, *F*_*AIZ*_) – and specifically, it will involve the part of one or more of those trios with strong two-species feedback (*F*_*SI*_, *F*_*SZ*_, and/or *F*_*IZ*_, respectively; Fig. 1D).

Finally, the fourth level of feedback (*F*_*4*_) captures loops through the whole resource-host-parasite system. *F*_*4*_ plays two roles in generating instability. First, as forecast above, if *F*_4_’s strength exceeds that of the lower levels (as combined in *HC*; eq. 7), then oscillations can arise. The longer, slower feedback loops in *F*_4_, when dominant, can trigger delays that provoke oscillations. Second, when *F*_4_ is positive at an ‘interior equilibrium’ (one with positive densities), that equilibrium is unstable and separates alternative stable states. To evaluate the role of resources at this level, we derived and organized *F*_4_ from a resource-centered perspective that sums four components. The ‘*via resource*’ (green box, Fig. 1E) component multiplies the effect of the resource on itself (*J*_*AA*_) times feedback in the epidemiological subsystem (-*F*_*SIZ*_). Resource self-facilitation (*J*_*AA*_ > 0) could influence oscillations (*HC* < 0) or alternative states (*F*_4_ > 0) only through this part of *F*_4_. Then, in the ‘*via susceptibles*’ (blue box), the resource affects susceptible hosts (*S*), and then *S* responds back in a four species net effect (*N*_*AS*.*IZ*_) composed of a snap back to *A*, a path then to *I* first and back to *A*, then finally a path to Z and then back through *I* to *A*. Those first two components (*via resources, via susceptible hosts*) appear in all three model variations. The ‘*via infected hosts*’ component (red box) is introduced by type III-clearance based exposure (variants 1 and 3) given its dependence on *J*_*IA*_. The resource affects *I*, then *I* returns feedback to *A* in three different ways (direct, via *S* first, or via *Z* first). Finally, the ‘*via propagules*’ component (grey box) enters due to resource-dependent propagule yield (*J*_*ZA*_) in variation 2. Assuming *J*_*ZA*_ > 0, the resource-propagule-susceptible host paths (direct and indirect) both increase resources by reducing consumption from *S* and *I*. Hence, these two ‘cascade fueling’ (*CF*) loops produce positive feedback. In contrast, the direct and indirect resource-propagule-infected hosts paths both depress resources by consumption. Hence, these two ‘consumption constraint’ (*CC*) loops produce negative feedback. Importantly, when strong enough, the *CF* loops introduce positive feedback that can turn *F*_4_ > 0.

## RESULTS: RESOURCES, POSITIVE FEEDBACK, AND ‘LOOP TRACING’

### Overview of model behavior and bifurcations (Fig. A1)

All model variants were studied with the aid of Mathematica and the Dynamica package for 2D bifurcation diagrams. All code used to generate these analyses and results is publicly available at [*redacted for double blind peer review*]. To examine the qualitative behaviors of these model variants, we varied enrichment (carrying capacity of the resource, *K*) and background death rate of susceptible hosts (*d*_*S*_). Bifurcation diagrams for each variant (1-3) show the qualitative behaviors that each variant produced in *K*-*d*_*S*_ space (summarized in Fig. A1; also Fig. 2A, 4A, 5A). The first, *S*_*in*_, depicts when the host (*S*) can invade the resource-only *A* equilibrium. The second, *R*_0_ = 1 (red), shows when the parasite (*I* and *Z*) can successfully invade the *AS* equilibrium given its net reproductive ratio (*R*_0_). Both *S*_*in*_ and *R*_0_ = 1 are ‘transcritical’ bifurcations. Third, Hopf bifurcations (dashed [*AS*] and solid [*ASIZ*] gray lines) denote the onset of oscillations, transitioning from a stable point to a stable limit cycle (yellow regions, *ASIZ*). Fourth, fold bifurcations (solid black lines) denote the introduction of a stable and an unstable equilibrium, marking the onset of alternative states (solid purple areas). Lastly, a limit point cycle (dashed blue lines) combines these two phenomena, introducing both a stable and an unstable limit cycle (like a fold for cycles). The unstable cycle separates the new stable cycle from an existing stable equilibrium. As shown below, these five bifurcations divide up carrying capacity (*K*) - mortality (*d*_*S*_) space differently for each model variant. We used loop analysis to demonstrate how resources drove Hopf and fold bifurcations, and the corresponding changes in model behavior, for each variant. Tracing the role of resources via feedback loops generates the fresh biological insights that lie at the heart of this paper.

**Figure 2:**
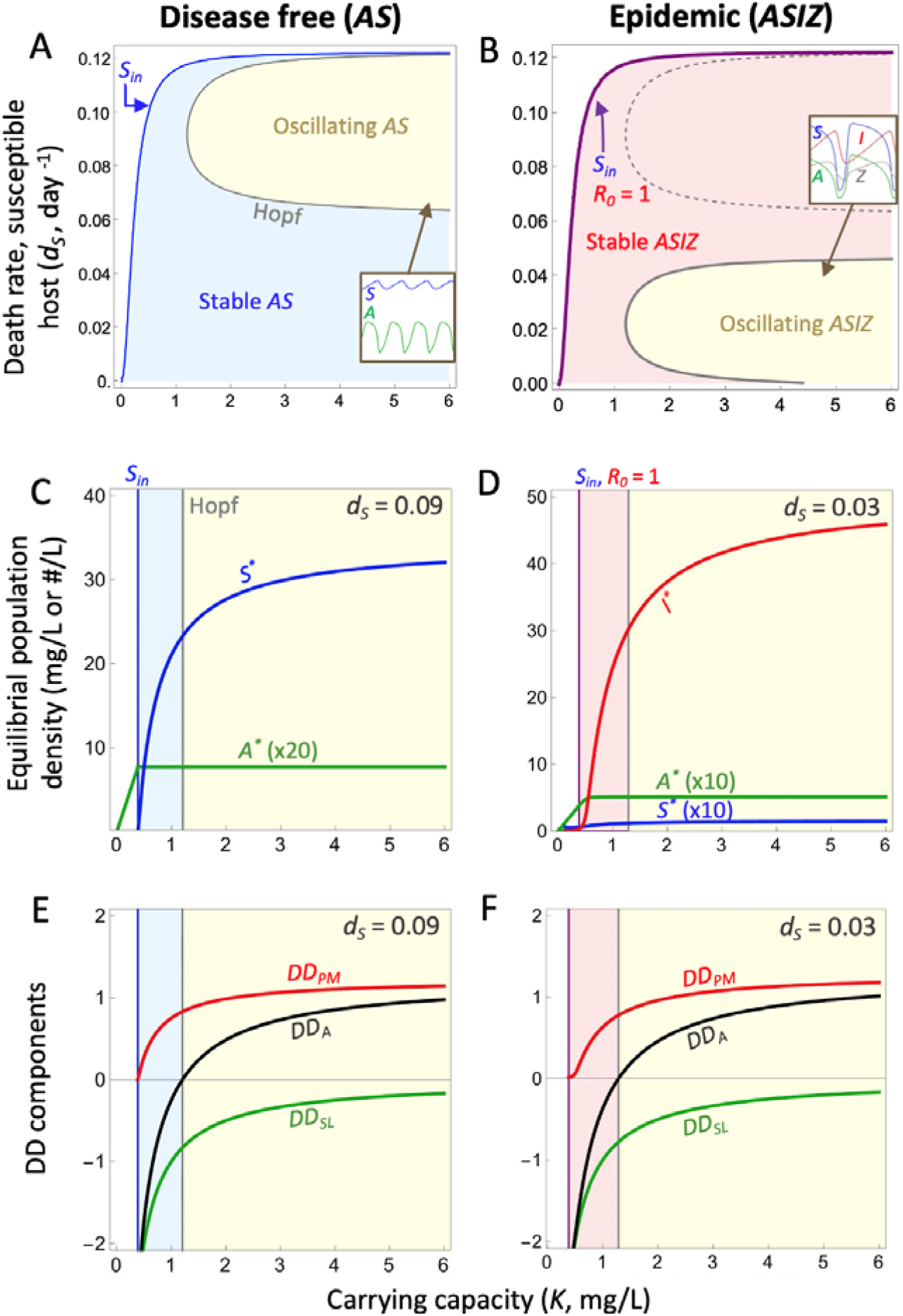
Positive density-dependence of the resource and oscillations in disease-free and epidemic systems. **(A)** Without disease (*AS*), a type III functional response of the grazer-host generates resource-host oscillations (yellow) at high carrying capacities of the resource (*K*) and death rates of hosts (*d*_*S*_). Inset: oscillating *AS* dynamics. **(B)** In the epidemic model (*ASIZ*), type III clearance also generates oscillations at similar carrying capacity (*K*) as the disease-free (*AS*) subsystem (dashed gray line), but at lower death rate of susceptible hosts (*d*_*S*_). Inset: oscillating *ASIZ* dynamics. Densities of *A*^*\**^ and *S*^*\**^ with increasing *K* for the **(C)** disease-free and **(D)** epidemic system. Equilibria are plotted in the oscillation region, not cycle averages, as feedback is calculated on equilibria. Net density-dependence of the resource (*DD*_*A*_; black line) and its components from predation mortality (red: *DD*_*PM*_) and self-limitation (green: *DD*_*SL*_) along gradients of *K* for the **(E)** disease-free and **(F)** epidemic system. When *DD*_*A*_ > 0, the *AS* system oscillates, but the border for epidemic oscillations is not exactly at *DD*_*A*_ = 0. (Disease-free: *d*_*S*_ = 0.09, epidemic *d*_*S*_ = 0.03; see also Table 1).

### Variant 1: How type III clearance creates non-epidemic and epidemic oscillations (Fig. 2)

Type III clearance creates oscillations in disease-free and epidemic systems by introducing positive density-dependence, hence self-facilitation, for the algal resource. The disease-free (*AS*) system produces two behaviors in carrying capacity (*K*)-death rate (*d*_*S*_) space (Fig. 1A): hosts invade past *S*_*in*_, then oscillations start in the region enveloped by the Hopf bifurcation at relatively high *K* and *d*_*S*_ (Fig. 2A inset). The epidemic (*ASIZ*) model produces similar behavior (Fig. 2B): hosts invade past *S*_*in*_, the parasite invades when net reproductive ratio *R*_0_ > 1 (Diekmann et al. 2010, calculated on *AS* cycles in oscillating region [Bate and Hilker 2013], see Appendix section 2, Fig. A2). Oscillations start in the region enveloped by the Hopf bifurcation (Fig. 1B and inset). Oscillations occur at significantly lower death rates *d*_*S*_ in the epidemic model than in the disease-free model (see also Appendix section 2).

In both the disease-free (*AS*) and epidemic (*ASIZ*) models, positive density-dependence of the resource (*DD*_*A*_) from consumption-based mortality trigger oscillations directly or indirectly via feedback loops. The location of oscillations in *K-d*_*S*_ space shifts due to differential responses of resource, host, and propagule densities and their influence on *DD*_*A*_. In the *AS* model, resources sit at the host’s minimal resource requirement (*A*_*S*_*), unchanging with *K*, while host density, *S*\*, increases with *K* (Fig. 3C). Similarly, in the *ASIZ* model, resource density (*A*) increases until host invade (at *S*_*in*_, where *K* = *A*_*S*_*). Almost immediately thereafter (given parameters), the disease subsystem (*I* and *Z*) invades when its net reproductive ratio *R*_0_ > 1. Once the parasite invades, total host population (*H*^*\**^ *= S*^*\**^ *+ I*^*\**^) increases with high infection prevalence while resources rise to the host’s updated minimal requirement (*A*_*H*_*; Fig. 2D). Given these similar density responses, *DD*_*A*_ (hence *J*_*AA*_) flips positive with enrichment (*K*) because the self-limitation component (*DD*_*SL*_) weakens while the predation mortality component (*DD*_*PM*_) strengthens (becomes more positive; Fig. 2E,F). As shown elsewhere (Appendix Section 2, Fig. A3), the region of oscillations and positive density-dependence of the resource shift from higher background mortality (*d*_*S*_) without parasites to lower *d*_*S*_ with parasites. These shifts arise because the epidemic shifts densities, particularly resources, due to a parasite-mediated trophic cascade.

**Figure 3:**
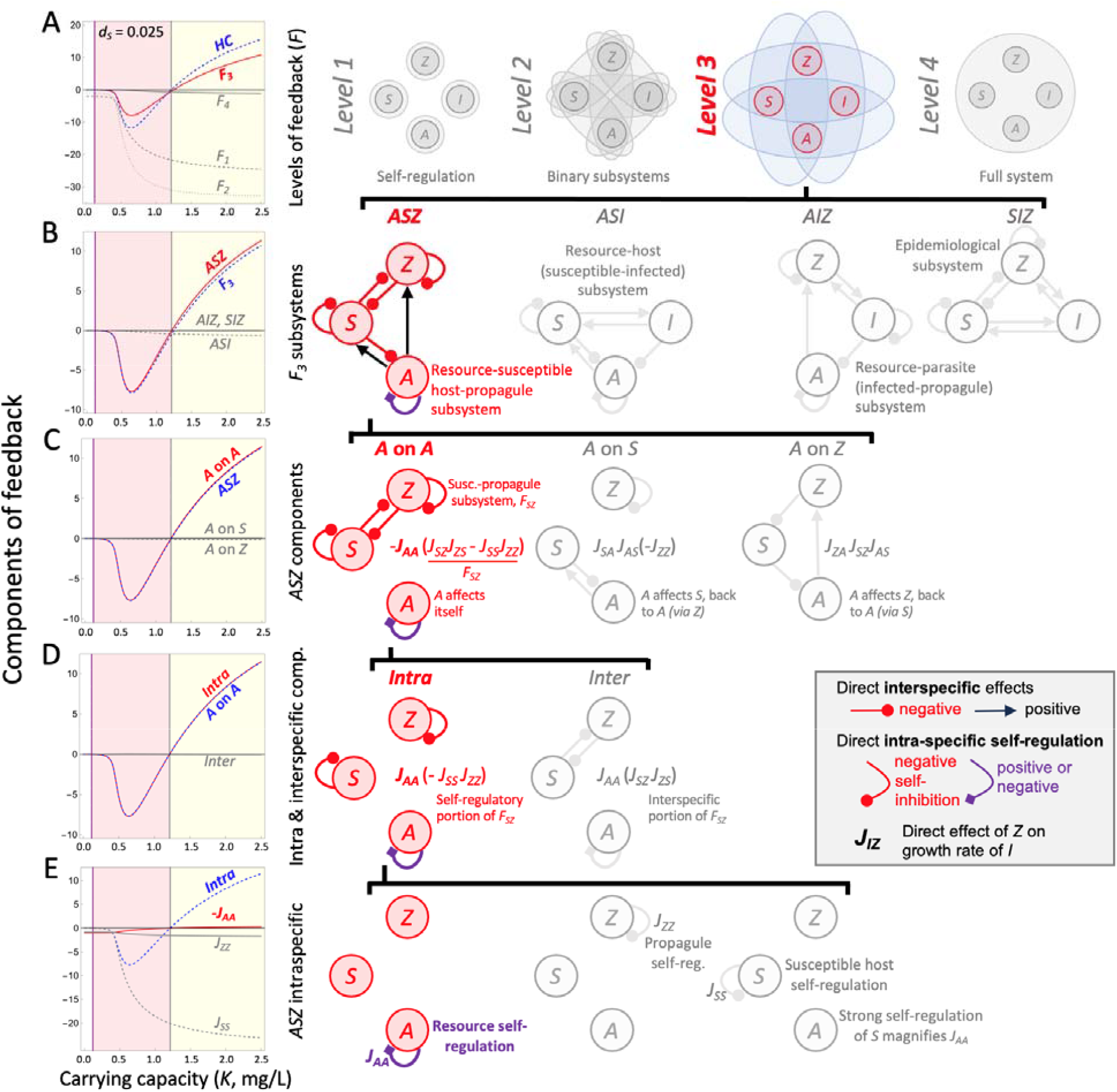
‘Loop tracing’ the genesis of epidemic oscillations in variant 1 of a resource-host-parasite model (ASIZ). In ‘loop tracing’, the focal component (blue) is shown with the responsible subcomponent (red) vs. less relevant ones (gray; left column), with accompanying loop diagrams (right). **(A)** Epidemic oscillations start when the Hopf criterion (*HC*, eq. 7) becomes positive. At this point, feedback in the entire system (*F*_4_) exceeds that in the lower levels (*F*_1_, *F*_2_, *F*_3_) combined. The flip in sign of *HC* most closely correlates with that of level three feedback, *F*_3_. **(B)** The resource-susceptible host-propagule (*ASZ*) subsystem primarily drives *F*_3_. **(C)** Within the *ASZ* subsystem, the ‘*A* on *A*’ component shapes *F*_*ASZ*_ more than ‘*A* on *S*’ and ‘*A* on *Z*’ components. **(D)** That ‘*A* on *A*’ component involves *J*_*AA*_ times feedback in the two-species *SZ* subsystem (*F*_*SZ*_ = *J*_*SZ*_ *J*_*ZS*_ – *J*_*SS*_ *J*_*ZZ*_). The intraspecific term, -*J*_*SS*_ *J*_*ZZ*_, changes with ‘*A* on *A*’. **(E)** In the intraspecific term, *J*_*AA*_ is smallest but shifts sign with the onset of oscillations (see also Table 1, Fig. A4).

Despite quantitative differences in the onset of oscillations and positive density-dependence of the resources, oscillations always began once or near where *DD*_*A*_ becomes positive. In the resource-host alone (*AS*) model, oscillations commence exactly once *DD*_*A*_, hence *J*_*AA*_, become positive. With only two species, oscillations begin when level 1 feedback (*F*_1_) becomes positive (Fig. 1B; Puccia and Levins 1985). Because *F*_1_ = *J*_*AA*_ (as hosts experience no self-regulation [*J*_*SS*_ = 0]), once resources self-facilitate, the system oscillates – the connection from resource ecology to oscillations is straightforward. However, as described above, oscillations with four ‘species’ (dimensions) start when the top level of feedback (*F*_4_) exceeds strength of feedback in the lower levels (*F*_1_, *F*_2_, *F*_3_) as arranged in the Hopf criterium (*HC*; eq. 7). Hence, self-facilitation of the resource – if it indeed underlies oscillations during epidemics – must act by weakening the lower levels of feedback vs. the topmost. Therefore, as forecast above, *J*_*AA*_ > 0 could weaken *F*_1_, some ‘resource-dependent’ components of *F*_2_, and some of the three *F*_3_ subsystems containing resources (*ASI, ASZ, AIZ*).

To test the hypothesis that resource self-facilitation still underlies oscillations in the resource-host-parasite system, we ‘trace’ the effect of *J*_*AA*_ through feedback loops in variant 1. We start at the largest scale: near where the Hopf criterium (*HC*) flips positive to start oscillations (at *HC* = 0), the third level of feedback, *F*_3_, notably changes from negative to positive (Fig. 3A). That observation implicates *F*_3_’s role in generating instability, as *F*_1_ and *F*_2_ strengthen stabilizing feedback (i.e., become more negative). Then, within *F*_3_, feedback in the resource-susceptible host-propagule subsystem (*ASZ*) most closely mirrors *F*_3_. The other two subsystems with resources (*ASI, AIZ*) and the epidemiological subsystem (*SIZ*) do little to counter *ASZ*’s influence. Further tracing, now within the *ASZ* subsystem, reveals that the component containing *J*_*AA*_ drives *F*_*ASZ*_ (*J*_*AA*_ [-*F*_*SZ*_]; Fig. 3C). Feedback in the *SZ* binary subsystem (*F*_*SZ*_) can be separated into intraspecific (*J*_*SS*_ *J*_*ZZ*_) and interspecific components (*J*_*SZ*_ *J*_*ZS*_), both scaled by *J*_*AA*_. Only the intraspecific components, *J*_*AA*_ *J*_*SS*_ *J*_*ZZ*_, shift from negative to positive near the Hopf (Fig. 3D). Finally, within the intraspecific components, only *J*_*AA*_ flips sign near the Hopf; both *J*_*SS*_ and *J*_*ZZ*_, while larger in magnitude, become more negative at that transition (Fig. 3E). Consequently, oscillations begin just about when the resource experiences positive density-dependence (*DD*_*A*_ > 0), hence self-facilitation (*J*_*AA*_ > 0). However, *J*_*AA*_ works indirectly, by weakening level three feedback, in a subsystem with susceptible host and propagules, because each experience strong self-regulation themselves.

#### Summary of ‘loop tracing’ of oscillations in variant 1

As in the disease-free (*AS*) system, positive density-dependence of the resource (*DD*_*A*_), hence self-facilitation (*J*_*AA*_), created by mortality imposed by the host, primarily triggers oscillations in the epidemic (*ASIZ*) model. However, in four dimensions, intraspecific, direct positive feedback (*J*_*AA*_ > 0) cannot work alone like it can in two dimensions. Instead, building from the bottom of our ‘loop tracing’ up: the switch to resource self-facilitation (*J*_*AA*_ > 0) is magnified by intense self-limitation of susceptible hosts (in *J*_*SS*_) but also self-limitation of propagules (*J*_*ZZ*_). Because *J*_*SS*_ *J*_*ZZ*_ together are so strong, self-facilitation of the resource creates strong feedback in the *ASZ* subsystem. (In the other two subsystems with resources [*ASI, AIZ*], binary feedbacks in the epidemiological subsystems [*F*_*SI*_ and *F*_*IZ*_, respectively] were too weak to magnify *J*_*AA*_ [Fig. A4]). Hence, *J*_*AA*_ weakens negative feedback in *F*_*ASZ*_, which eventually turns *F*_3_ positive. Weakening *F*_3_ then ensures that the combination of faster, shorter, lower levels of feedback (*F*_1_-*F*_3_, combined in the *HC*) is outweighed by feedback in the longer, slower loops of total system feedback (*F*_4_). Once feedback in *F*_4_ predominates, delays in the system become strong enough to commence epidemic oscillations (see Appendix section 4 for more details). Nonetheless, through loop tracing, we still found that positive density-dependence of the resource, hence resource self-facilitation, indirectly created epidemic oscillations.

### Variant 2: Alternative stable states with resource-dependent yield of propagules (Fig. 4)

In variant 2, resources create a different form of instability through a different mechanism of positive feedback. Variant 2 assumes linear clearance rate (eq. 2A) and resource-dependent propagule yield (eq. 5B). As shown above (Fig. 1E), resource-dependence of propagule yield introduces a new, positive, interspecific, direct link between resources and propagules (*J*_*ZA*_) into parts of both level three (*F*_3_) and level four feedback (*F*_4_, in the ‘*via propagules*’ component). However, the genesis of alternative stable states and resulting Allee effects for the parasite involve *F*_4_, total system feedback. Specifically, alternative stable states arise when *F*_4_ becomes positive. As forecasted above, *F*_4_ can become positive when the injection of *J*_*ZA*_ adds cascade-fueling (*CF*) loops that generate positive feedback. If *CF* loops exceed negative feedback in the new consumption constraint (*CC*) and other basal loops (‘*via resources*’ + ‘*via susceptible hosts*’; Fig. 1E), then total system feedback can become positive (Appendix section 4).

**Figure 4:**
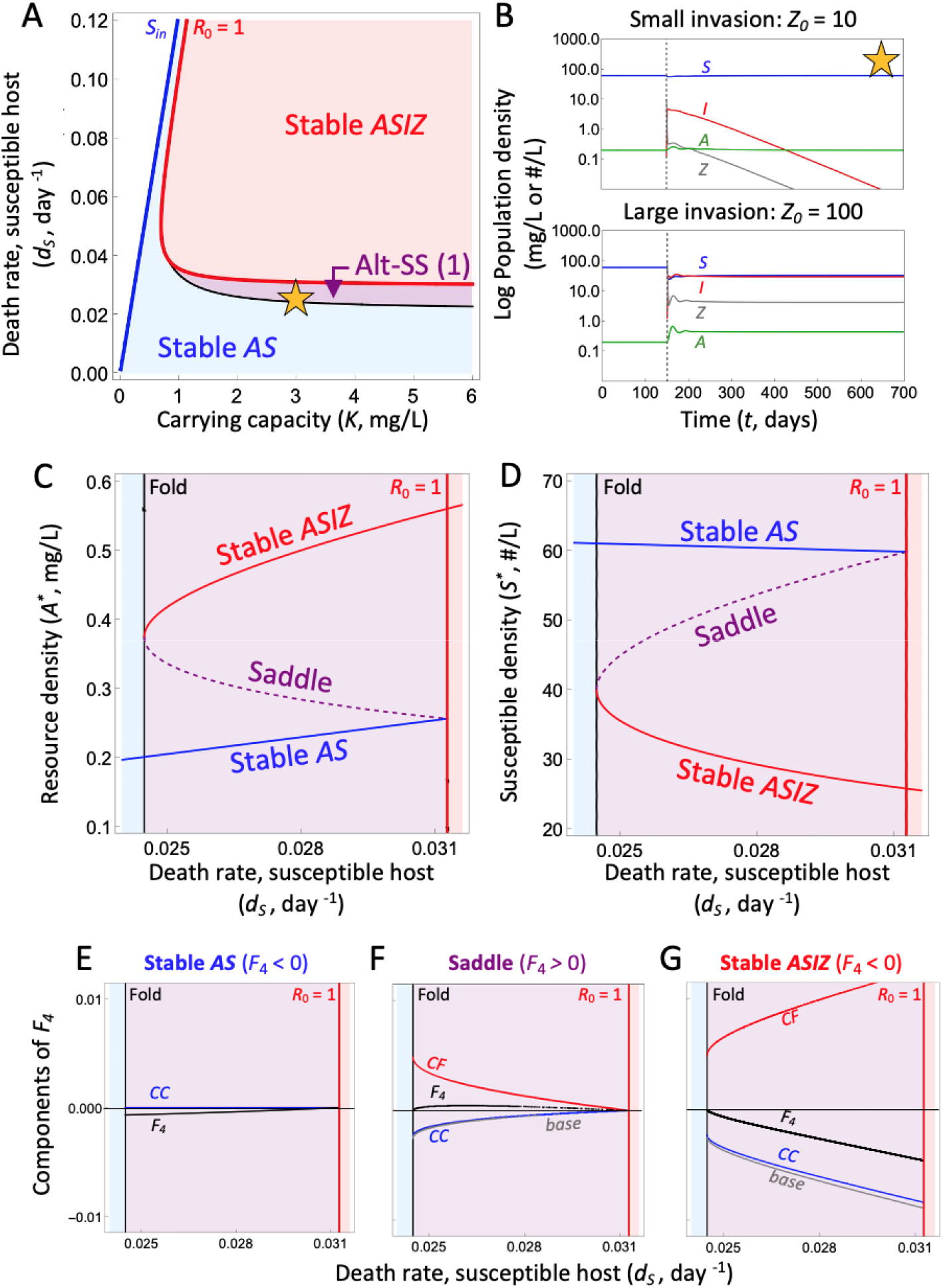
Genesis of alternative stable states and Allee effects in variant 2 of a resource-host-parasite model. **(A)** Resource-dependent propagule yield, σ(*A*), generates bi-stability in a narrow band of host death rates (purple region, ‘Alt-SS (1)’). **(B)** A small introduction of parasites at day 150 (*Z*_0_ = 10) cannot start the epidemic (top), but a larger initial invasion (*Z*_0_ = 100) can (at yellow star *K*-*d*_*S*_). Equilibrium densities of **(C)** resources (*A*^*\**^) and **(D)** susceptible hosts (*S*^*\**^) with increasing death rate (*K* = 3). Alternative stable states (shaded purple) sit between a fold (lower *d*_*S*_) and *R*_0_ = 1. An intermediate saddle (purple dashed curve) separates a stable *AS* (blue) from a stable *ASIZ* (red) equilibrium. **(E-G)** Via the interspecific *J*_*ZA*_ term, resource-dependent production of propagules introduces positive, destabilizing feedback from ‘cascade fueling’ (*CF*) loops and negative, stabilizing feedback from a ‘consumption constraint’ (*CC*) to ‘base’ level four feedback (Fig. 1E). These *CF* (+, red), *CC* (-, blue), and ‘base’ (-, grey) loops combine into *F*_4_ (black) for the **(E)** stable disease-free (*AS*), **(F)** unstable saddle (*ASIZ*), and **(G)** stable (*ASIZ*) case (see also Table 1).

To validate this argument, we dissect the region of alternative stable states in variant 2. This region (‘Alt-SS(1)’, purple) sits sandwiched between parasite invasion when rare (*R*_0_ = 1, red curve, a transcritical bifurcation) and a fold bifurcation (black curve, Fig. 4A). In this region, relatively low background death of susceptible hosts (*d*_*S*_) keeps resource density small (*A*_*S*_*) – too low to allow a small invasion of propagules to infect enough hosts to produce enough propagules to propel the epidemic forward (so, *R*_0_ < 1 here; Fig. 4B top). In contrast, a sufficiently large invasion (e.g., *Z*_0_ = 100; bottom panel, Fig. 4B) triggers a trophic cascade that increases resources (*A*) that then fuels enough production of propagules to propel the epidemic onward. Hence, the parasite in this region experiences an ‘Allee effect’, where parasite invasion must push the system beyond a threshold for epidemics to commence. The fold bifurcation introduces both a stable epidemic (*ASIZ*) equilibrium (*F*_4_ < 0) and an unstable one (a saddle: *F*_4_ > 0; Fig. 4C,D). This saddle separates the stable epidemic from the stable disease-free equilibrium and shows the threshold that determines the Allee effect (high enough resource density [Fig. 4C] caused by a sufficient plunge in susceptible hosts [Fig. 4D]). Once *d*_*S*_ rises past the *R*_0_ = 1 threshold, only the stable disease equilibrium exists as the parasite can invade when rare.

The strength of negative vs. positive feedback at three relevant equilibria – the two stable and one saddle – governs the Allee effect. Neither positive feedback from ‘cascade fueling’ (*CF*) loops nor negative feedback from their companion ‘consumption constraint’ (*CC*) loops operate at the stable, disease-free, host-resource (*AS*) equilibrium (Fig. 4E). There, that *AS* equilibrium provides net negative feedback (*F*_4_ < 0). Then, the relative strength of positive cascade fueling (*CF*) and negative consumption constraining (*CC*) plus ‘base’ feedback loops (‘*via resources*’ and ‘*via susceptible hosts*’ [Fig. 1E) determine stability of the two epidemic equilibria introduced by the fold. At the intermediate saddle separating the alternative stable states, positive feedback from *CF* exceeds negative feedback from *CC* plus the ‘base’ (grey) loops (hence *F*_4_ > 0; Fig. 4F). Conversely, at the stable disease (*ASIZ*) equilibrium, negative feedback from *CC* and base loops overpowers the positive *CF* loops (hence *F*_4_ < 0; Fig. 4G). Hence, the positive, direct, interspecific connections between resources and propagules introduced by resource-dependent propagule yield (*J*_*ZA*_ > 0) indeed do create alternative stable states and Allee effects of the parasite. Using ‘loop tracing’, we pinpoint why: *J*_*ZA*_ introduces ‘cascade fueling’ loops that generate positive feedback by reducing resource consumption from hosts.

#### Comparing variants 1 and 2

The basal resource, *A*, lies at the center of two different forms of positive feedback that create two forms of instability. In variant 1 (type III clearance rate, constant propagule yield), the disease (*ASIZ*) system could oscillate. Review of the well-known disease-free (*AS*) system showed how positive, individual-level density-dependence of the resource (*DD*_*A*_ > 0) and corresponding population-level self-facilitation (*J*_*AA*_ > 0) directly trigger these oscillations. In the disease system (variant 1), *J*_*AA*_ > 0 strongly influenced their onset, too. However, in that case *J*_*AA*_ generated oscillations by weakening a combination of faster, shorter lower-level feedback (*F*_1_, *F*_2_, and *F*_3_) vs. slower, longer upper-level feedback (*F*_4_, as arranged in the Hopf criterion [*HC*]). We traced *J*_*AA*_’s role in particular loops (esp. the *ASZ* subsystem of *F*_3_) to make that argument. In contrast, in variant 2 (linear clearance rate, resource-dependent propagule yield) we saw how resources generated positive feedback at level 4 (*F*_4_). This time, the intraspecific effect (*J*_*AA*_) did not feature. Instead, the interspecific effect of *A* on *Z* (in *J*_*ZA*_, introduced by resource-dependent production of propagules) created ‘cascade fueling’ loops that generated positive feedback in *F*_4_. When the magnitude of *CF* loops exceeded the sum their partner consumption constraint (*CC*) loops and the other basal *F*_4_ loops, positive feedback ensured one of the disease equilibria was a saddle that separated the stable disease-free from the stable disease states (thereby creating Allee effects for the parasite).

### Variant 3: Instabilities with type III clearance & resource-dependent propagule yield (Figs. 5, A3)

Variant 3 of the resource-host-parasite model combines resource-dependence of feeding / exposure rate (type III clearance; eq. 2B) with resource-dependent yield of propagules (eq. 5B). Hence, we hypothesized that it would feature both forms of resource-driven instability. However, the feedback loops underlying those instabilities have now become the most complex (Fig. 1). In particular, total system feedback (*F*_4_) now includes ‘*via propagule*’ loops introduced by the direct effect of resources on propagules (*J*_*ZA*_, usually positive) and ‘*via infected*’ loops added by the direct effect of resources on infected hosts via exposure (*J*_*IA*_; Fig 1E). Despite different and more complicated loop structures in this variant, the model still produces simple epidemic oscillations (in the ‘oscillating *ASIZ*’ region; Fig. 5A). And once again, loop tracing revealed that self-facilitation of the resource (*J*_*AA*_ > 0), introduced by grazing mortality (and *DD*_*A*_ > 0), triggered them by weakening lower levels of feedback (*F*_1_, *F*_2_, *F*_3_) relative to the total system feedback (*F*_4_). However, the details differ this time (Appendix section 4, Fig. A5). Now *J*_*AA*_ works by weakening feedback at level 1 (*F*_1_) – self-inhibition of the resource weakens and counters increasing self-inhibition of the infected hosts (*J*_*II*_). Additionally, *J*_*AA*_ weakens level two (*F*_2_) mostly by weakening feedback in the resource-propagule subsystem (*F*_*AZ*_; see Fig. A5 for details). Nonetheless, the same basic outcome arises. Epidemic oscillations start because density-dependence of the resource (*DD*_*A*_) creates self-facilitation (*J*_*AA*_) that weakens stabilizing lower levels of feedback relative to total system feedback.

**Figure 5:**
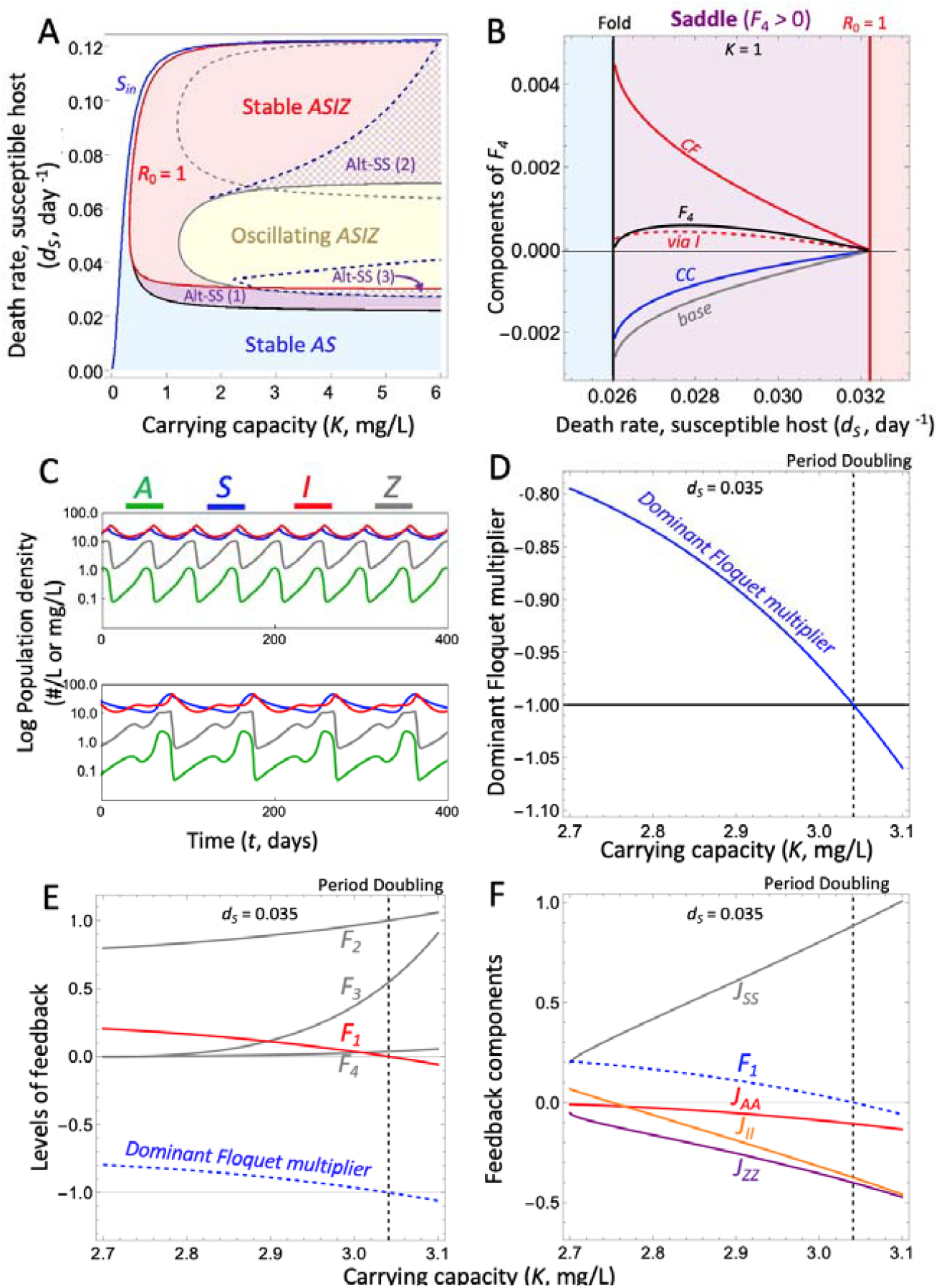
Alternative stable states and period doubling with resource-dependent exposure and propagule yield. **(A)** With increasing carrying capacity of the resource (*K*) and host death rate (*d*_*S*_), variant 3 produces simple oscillations (like in variant 1) and bi-stability (Alt-SS [1], like in variant 2). Two new regions of alternative states also emerge (Alt-SS [2] and Alt-SS [3]; see also Appendix section 5). **(B)** Like variant 2 (Fig. 4), alternative stable states in variant 3 are driven by cascade-fueling (*CF*; red line) loops. However, the addition of *via infected hosts* (‘via *I*’) loops (dashed red line) from type III clearance further contributes to positive feedback in *F*_*4*_ underlying Alt-SS (1). Color conventions follow Fig. 4. **(C)** A period-doubling route to chaos emerges with *K* (at *d*_*S*_ = 0.035). Top (*K* = 2.5): simple oscillations. Bottom (*K* = 3.5): period-doubled cycles with alternating small and large epidemics. **(D)** The dominant Floquet multiplier crosses -1 at the period-doubling bifurcation (here ∼*K* = 3.041). **(E)** Strengthening level 1 feedback *F*_*1*_ undergoes a sign change when the period doubling bifurcation (dashed vertical line) occurs. **(F)** Strengthening *J*_*AA*_, (red), *J*_*II*_ (orange), and *J*_*ZZ*_ (purple) drive the change in *F*_*1*_ that yields the period-doubling bifurcation.

In this variant, simple alternative stable states also create Allee effects for the parasite. Like in variant 2, these Allee effects in the Alt SS-1 region (Fig. 5A) are caused by the introduction, by a fold, of an unstable (saddle) epidemic equilibrium with positive total feedback (*F*_4_ > 0). That saddle separates stable disease-free equilibrium (*AS*) and a stable epidemic equilibrium (*ASIZ*). Like in variant 2, the saddle has positive total feedback because positive, direct resource-propagule links (*J*_*ZA*_) introduce ‘cascade fueling’ loops (Figs. 1E,5B). Those positive *CF* loops exceed the strength of negative consumption constraint (*CC*) loops also introduced by *J*_*ZA*_. However, whole system feedback also includes positive feedback introduced by direct resource-infected links (*J*_*IA*_ > 0). Positive feedback in the ‘*via infected*’ loops arises because higher exposure increased infected hosts, thereby increasing propagules which remove susceptible hosts, reducing resource consumption (in the third loop of the ‘*via infected*’ component of *F*_4_; the other two produced negative feedback; Fig. 1E). Nonetheless, this contribution to positive feedback from *J*_*IA*_ remains smaller relative to the *CF* loops (Fig. 5B).

Additionally, the insertion of non-linear, type III clearance rate into resource mortality, exposure, and propagule yield terms produces several new and more exotic regimes into variant 3 (Fig. 5A). For instance, two new types of alternative states are introduced by limit point cycle (LPC) bifurcations (fold bifurcations of cycles, birthing both an unstable and a stable orbit). At high carrying capacity of the resource (*K*) and high death rate of susceptible hosts (*d*_*S*_), a second region of alternative stable states (Alt-SS [2]) exhibits epidemics at either a stable equilibrium or a stable limit cycle (oscillations; Fig. 5A). An unstable cycle separates them. Then, at lower *d*_*S*_ and near the simpler Allee effect region (Alt-SS[1]), a third type of alternative states, Alt-SS (3), arises where Alt-SS (1) and epidemic oscillations collide (Fig. 5A), producing either stable host-resource (*AS*) or oscillatory disease (*ASIZ*) dynamics, depending on initial conditions – variation on the Allee effect of Alt-SS (1). Finally, the oscillating disease (*ASIZ*) system in the Alt-SS (3) region can undergo a period doubling route to chaos with increasing *K* (Fig. 5A – for evidence of chaos, see Appendix Section 5, Fig. A6).

Unfortunately, because loop analysis was designed for points, not orbits, we cannot readily ‘trace’ feedback behind these destabilized cycles with known methodology. However, we would like to suggest – as a call for future work – that loop tracing might explain the biology underlying the destabilization of cycles in a period doubling example in Alt-SS (3). Biologically, that period doubling introduces alternating peaks of smaller vs. larger epidemics (indexed by infected hosts [*I*] in Fig. 5C). However, to find where that doubling arises and why, the approach above needs tweaking. First, we calculate feedback loops on the *ASIZ* system’s ‘monodromy’ matrix, a cousin of a Jacobian matrix described elsewhere (Appendix section 5). Then, to find the first period doubling, we calculated Floquet multipliers (an analogue to eigenvalues of a Jacobian: Klausmeier 2008). If all real parts of Floquet multipliers are between -1 and 1, the orbit is stable. Period doubling happens once the dominant Floquet multiplier falls below -1 (e.g., at ∼*K* = 3.041 in the example provided; Fig. 5D).

Once we found this border, we hypothesized that it would correspond with feedback calculated over an epidemic cycle. We observed a sign change in level 1 feedback (*F*_*1*_), exactly where the period doubling bifurcation occurs (Fig. 5E). The other levels of feedback (also calculated over the cycle) did not change in a comparable way (Fig. 5E). As the cycle destabilizes (period-doubles), *F*_1_ is strengthening (becoming more negative). Dissecting *F*_1_ further, self-inhibition of the resource (*J*_*AA*_), infected hosts (*J*_*II*_), and propagules (*J*_*ZZ*_) all strengthen, enough to outweigh self-limitation of susceptible hosts (*J*_*SS*_) to push *F*_*1*_ negative (Fig. 5E). Thus, it seems that these self-regulation terms, as summed in *F*_*1*_, drive the first period doubling bifurcation in the *ASIZ* model. Notice, then, how destabilization of the cycle involves strengthening negative self-inhibition of each species. Future work can more rigorously connect cycling dynamics to self-regulation to the genesis of period-doubled cycles. For now, this lead promises that loop tracing might work for more complex dynamics and destabilization of cycles.

## DISCUSSION

In this paper we focused on a continued challenge for theoretical thinking in ecology, that balancing complexity and realism against tractability and understanding. The simplest two-species (two-dimension) models of species interactions provide powerful ecological insights gleaned from highly tractable mathematical and graphical analyses. Arguably, for those reasons, the Rosenzweig-MacArthur model of predator-prey oscillations and the Lotka-Volterra model of competition have made such powerful, long-lasting impacts. With three or more species, models become more realistic, but the core biological insights become harder to rigorously understand. It remains possible to describe the ‘what’ of higher dimension models (predictions given assumptions) but it becomes harder to extract out the ‘how’ and ‘why’ behind those predictions. We propose that analysis of feedback loops (Puccia and Levins 1985; Dambacher et al. 2003, Novak et al. 2016; Simons et al. 2022; Cortez 2024) provide an underutilized method to systematically address ‘how’ and ‘why’ questions. To illustrate, we considered how and why resources create instabilities in resource-host-parasite systems with four dimensions. Based on previous work, we knew that epidemic oscillations could arise when grazer-hosts feed according to non-linear (saturating) functional responses (see below). Based on the Rosenzweig-MacArthur model, we knew that ‘safety in numbers’ from those functional responses directly trigger oscillations in predator-prey models with two equations. But how and why would that biology create oscillations in more complex and interconnected scenarios with parasites? Previous work also has shown that resource-dependent yield of infectious propagules can yield Allee effects for parasite invasion (see below). But how and why should direct resource-propagule effects translate into whole-system positive feedback required for such Allee effects? We addressed both problems by ‘tracing’ the key parts of system feedback that create both instabilities. Such tracing in these meso-scale models revealed answers to ‘how’ and ‘why’ questions about unstable dynamics in resource-host-parasite systems.

In our first epidemic model, we used loop tracing to understand how and why resources created epidemic oscillations. Based on decades of previous work, we knew that positive density-dependence of resources (*DD*_*A*_) was likely involved. Positive *DD*_*A*_ arises when crowding (i.e., increasing density) lowers per capita mortality inflicted by hosts (‘safety in numbers’, Murdoch et al. 2003) despite fitness costs of logistic-based self-limitation. Fitness benefits of ‘safety in numbers’ surface at higher carrying capacity of the resource (as in the paradox of enrichment: Rosenzweig & MacArthur, 1963; Rosenzweig, 1971) and, as shown here, at high background consumer (host) death rate. We also knew that virulence from infection raises the host’s minimal resource requirement during epidemics, yielding oscillations at lower background mortality. Therefore, epidemics can stabilize unstable dynamics (like in Anderson and May, 1978; Hilker and Schmitz, 2008; Hurtado et al., 2014; Simon et al. 2022) but destabilize other stable ones (like in Hilker et al. 2009).

However, with loop tracing, we could pinpoint how and why positive *DD*_*A*_ generated oscillations during epidemics. In more than two dimensions, oscillations arose when chains of ecological relationships weakened more stabilizing, faster, lower level feedbacks (working through shorter one, two, and three species ‘loops’) relative to the longer, top level of feedback (working more slowly through all four species). When top level feedback dominated, the delays that it produced created oscillations (Lever et al. 2023). Via two examples, we illustrated how weaking density-dependence (hence self-regulation) of the resource weakened lower-level feedbacks. In variant 1 (with type III clearance rate of hosts), resource self-facilitation largely weakened part of level three feedback through its connection to strong self-inhibition of susceptible hosts and propagules. In variant 3 (adding resource-dependent propagule yield to assumptions of variant 1), resource self-regulation weakened feedback in level 1 and level 2 (in particular, part of *F*_2_ with resource-propagule links). Despite differences in details, in both cases we showed via ‘loop tracing’ how the ‘safety in numbers’ response of resources underpins oscillating dynamics. Hence, we showed how and why the simple lesson of the Rosenzweig-MacArthur model could still manifest in more complex and interconnected webs of interactions. Since loop tracing applies generally, we propose that it could be used to understand how and why epidemic oscillations interact with other factors such as competitors of hosts (Cáceres et al. 2014) and resources (e.g., inedible producers, Kretzschmar et al. 1993), immune competitors of hosts (aka, diluters: Strauss et al. 2018, Keesing and Ostfeld 2021), seasonality (Aron and Schwartz, 1984), and host behavior (Althouse and Hebert-Dufresne, 2014).

With loop tracing, we also showed that resources can generate alternative stable states, another form of instability, through another form of positive feedback. That positive feedback originates from resource-dependent yield of infectious propagules (Hall et al. 2009; Cressler et al. 2014; Fearon et al. 2025). An increase of yield with resources introduces a positive, direct, interspecific link between resources and propagules (called *J*_*ZA*_ here). Models with similar links show that the parasite can experience an Allee effect (Smith et al. 2015; Borer et al. 2023). Here, using loop tracing, we showed how and why positive feedback at a ‘saddle’ equilibrium creates the Allee effect. That saddle experiences positive total system feedback because *J*_*ZA*_ introduces positive ‘cascade fueling’ loops. In cascade fueling loops, propagules remove susceptible hosts which net increases resources (a cascade) that then fuels further production of propagules. These cascade-fueling loops exceed the strength those generating stabilizing, negative feedback. Hence, loop analysis pinpointed the causal, cascade-based mechanism underlying Allee effects. Variant 3, adding in type III functional responses and resource-dependent yield, also produced Allee effects. Once again, cascade fueling loops drove positive feedback. However, direct, positive links from resources to infected hosts (*J*_*IA*_ > 0) introduced positive feedback as well. Hence, loop tracing of this more complex case revealed two resource-disease links that create positive feedback loops underlying Allee effects. Application of this method could pinpoint how and why Allee effects arise in epidemics due to, e.g., predation (Hall et al., 2005; Ranjit et al., 2025) and in food webs more generally (Schröder et al. 2005; Scheffer, 2009).

The combination of positive feedback mechanisms from resource-dependent clearance/exposure rate and propagule yield also introduced more complicated behaviors. First, two new forms of alternative states emerged. One arose at higher background mortality of hosts; here, epidemics either oscillate or dampen to a stable equilibrium. The other new alternative state region lies adjacent to the original but separates the disease-free state from oscillating epidemics. In this region, more complex epidemic cycles arrived through ‘period doubling’, leading to chaos with enrichment (like in Bate and Hilker 2013; Saifuddin et al. 2016). In the simplest period-doubled case, epidemics cycle between low and high peaks of infected host density. Unfortunately, we do not know of established methods for apply loop tracing to orbits (cycles). However, based on preliminary analysis here, we suggest that maybe it could. First, we found where period doubling started (using known methodology with Floquet multipliers: Klausmeier et al. 2008). Then, we calculated feedback at all four levels over simple vs period-doubled cycles. With this loop tracing of the cycles, we found that feedback level 1 (*F*_*1*_), the sum of each species’ self-regulation, flipped sign right as first period doubling arose. However, the stable cycle destabilized as *F*_*1*_ strengthened (rather than weakened, as with oscillations). The increase in self-regulation resembles period doubling in the discrete time logistic model (Case 2000). This result suggests that, if further developed, loop tracing might apply to periodic, multi-periodic, and aperiodic dynamics that abound in ecology.

The loop tracing tools built here afford further development and testing of hypotheses regarding instabilities in disease dynamics. Five extensions immediately spring forward. First, other epidemiological components, such as immune clearance from infected or exposed classes, can depend on resources (Cressler et al. 2014; Cotter and Al Shareefi 2022). Additionally, foraging rate can depend on resource quality (Schatz and McCauley 2007; Penczykowski et al. 2014). Loop tracing can determine if those relationships change feedback loops that enhance or suppress instabilities in those systems. Second, parasites might introduce instability through other virulent effects on hists. For instance, strong castration (virulent fecundity reduction), a common parasite strategy (Hall et al. 2007, Lafferty and Kuris 2009), can provoke oscillations (Anderson and May 1978; Blower and Roughgarden 1987; Auld et al. 2014). With ‘loop tracing,’ one could test the hypothesis that weakening self-limitation of the epidemiological resource (susceptible hosts) underlies that instability. Third, heterogeneity of basal or host trophic levels might alter positive feedback. For instance, completely inedible producers or immune competitors of hosts might prevent the parasite-mediated cascades (as in Cáceres et al. 2014) or might stabilize cycles (as in McCann et al. 1998). Loop tracing might highlight how this action could sever, suppress, or enhance strength of resource-dependent loop connections that propagate or squelch positive feedback. Fourth, as mentioned above, further development of a ‘loop tracing of cycles’ might pinpoin why period doubling and other cycle bifurcations (e.g., limit point cycles) arsie. Finally, experimental manipulation of background death of hosts could shift epidemics from stable to various unstable dynamics (Fussman et al. 2000). It would be fascinating if parameterized loop tracing could quantitatively anticipate those shifts in behavior.

In the meantime, we show how and why resources can introduce positive feedback and destabilize disease dynamics. Resources can introduce positive feedback via their own ‘safety in numbers’ response to hosts that consume them. When acting at the population level, this self-facilitation cannot trigger epidemic oscillations alone; instead, it weakens the stabilizing effects of shorter, quicker loops relative to longer, slower ones. Alternatively, via resource-dependent production of propagules, resources can create positive feedback by linking key components within disease food webs. Those new connections can channel positive, total system feedback at equilibria that separate alternative states and create Allee effects. Both pathways involve the implications of resource release from parasite-mediated trophic cascades. When combined, resource-dependent exposure and yield can produce more complex, unstable dynamics (such as period doubling). Importantly, we linked biological relationships to positive feedback to dynamical outcomes by dissecting loops involving resources. In the future, similar dissections using loops could uncover deeper understanding and fresh insights into other stabilizing and destabilizing forces of disease and consumer-resource dynamics (Ramesh and Hall 2023, 2025). We hope that development of similar approaches can ‘trace’ the role of resources behind more complex behaviors as well.

## APPENDIX

This appendix provides additional support for the approaches and results in the main text. We first provide additional details regarding behavior of the disease-free system and the epidemic model variants (section 1). Then, we provide some additional mathematical detail for loop analysis and ‘tracing’ (section 2), results of the applications of loop tracing to the model variants (section 3), and full Jacobian matrices for each (section 4). Finally, we describe the analysis of the period doubling with an extension of loop tracing (section 5).

### Section 1: Full Jacobian matrices for all model variants

The loop analyses above depend on Jacobian matrices (**J**). These matrices describe the direct effects of each species on themselves (along the main diagonal) or on each other (Figs. 1A, A1A). We present them for the *AS* resource-host, disease free system and each variant of the *ASIZ* disease model. Some simplifications of derivatives of logistic production, *r*(*A*), feeding rate, *f*(*A*), and clearance rate, *c*(*A*) help simplify the matrices:

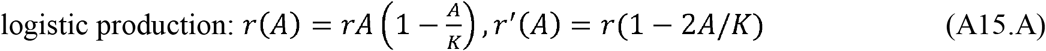

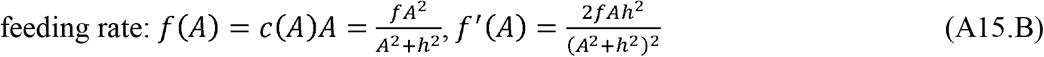

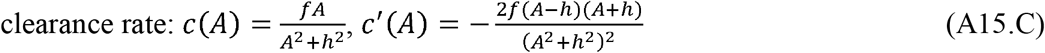

where the ′ notation denotes partial derivative (sensitivity) with respect to *A* (i.e., *r*□(*A*) = ∂*r*(*A*) / ∂*A*). The Jacobian for the *AS* case, **J**_**AS**_, is then (in order of *A*, then *S*):

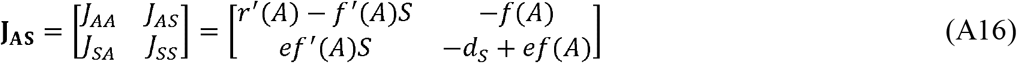

where element *J*_*XY*_ is the effect of an increase in density of *Y* on the growth rate of *X, G*_*X*_ (so *J*_*XY*_ = ∂*G*_*X*_/∂*Y*). For the three variations on the *ASIZ* disease model, the Jacobian (**J**_**ASIZ**_) has the general form, again (in order of *A, S, I*, and *Z*):

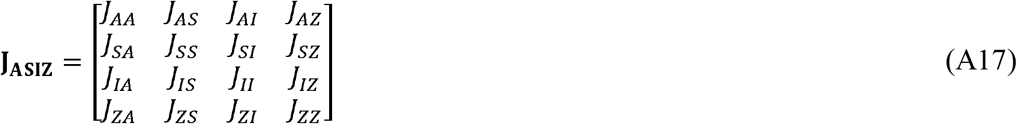

which, for variant 1 (type III clearance rate but fixed propagule yield), the Jacobian (**J**_**ASIZ**_) becomes:

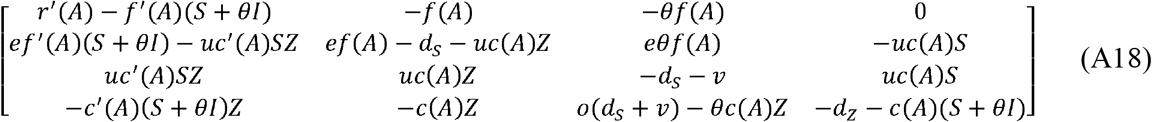

and for variant 2 (type I clearance and resource-dependent propagule yield) it is:

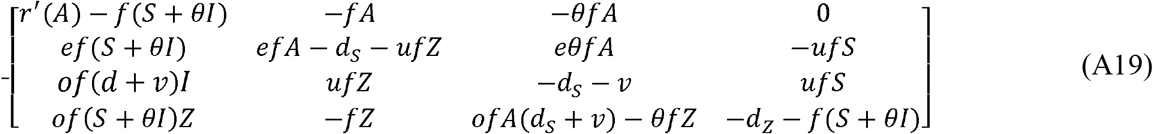

and for variant 3 (with type III clearance and resource-dependent propagule yield) it is:

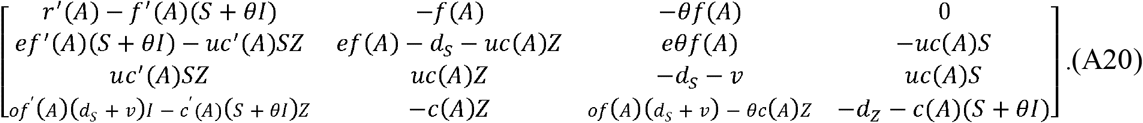

### Section 2: More analysis of the disease-free system and epidemic model variants

#### Behavior of the resource-host system

Type III clearance creates oscillations in disease-free and epidemic systems by introducing positive density-dependence for the algal resource. The disease-free (*AS*) subsystem is (with *G*_*j*_ for population growth rate and *r*_*j*_ for fitness):

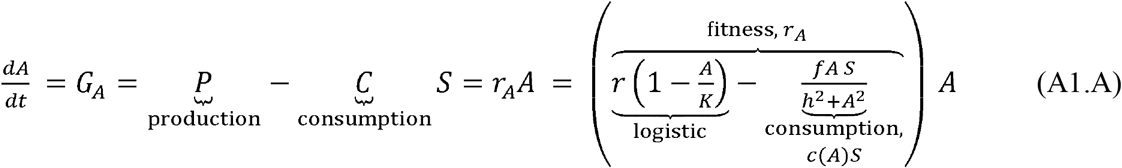

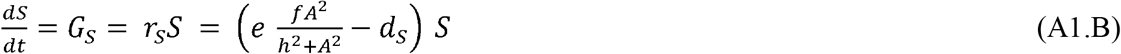

This *AS* disease-free model produces two behaviors in carrying capacity (*K*)-death rate (*d*_*S*_) space (Fig. 2A): hosts invade past *S*_*in*_, then oscillations start in the region enveloped by the Hopf bifurcation at relatively high *K* and *d*_*S*_ (Fig. 1A inset). Those oscillations follow a typical consumer-resource trajectory. Peak resource density (*A*) fuels peak host (*S*) density. High grazing pressure (peak *S* and intermediate *A*, hence intense per capita resource mortality) crashes the resource, and *S* declines through starvation. Without high consumption-based mortality and with little self-limitation, the low-density resource recovers, and the cycle renews.

Positive density-dependence of the resource from consumption-based mortality directly triggers oscillations in the disease-free system, following the same reasoning as the classic Rosenzweig-MacArthur model (Murdoch et al. 2003). The resource’s density-dependence (*DD*_*A*_) is the sensitivity of its fitness (*r*_*A*_; eq. A1) to an increase in its density (*A*):

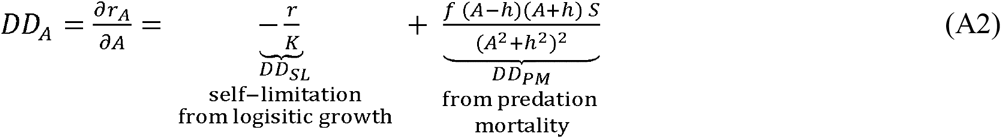

Negative density-dependence from the self-limitation term, *DD*_*SL*_ = -*r* / *K*, comes from the logistic portion of resource fitness (eq. A1). Without consumption (*A* only), *DD*_*SL*_ completely accounts for (negative) density-dependence of the resource; carrying capacity inhibits growth at high resource density. However, with type III clearance, hosts potentially introduce positive density-dependence, *DD*_*PM*_, through per capita mortality from consumption, *c*(*A*)*S*, a unimodal curve that peaks at the half-saturation constant (*h*, with peak clearance *f* / [2*h*]). ‘Safety’ (lower per capita mortality rate from consumption) comes from ‘rarity’ (*A* < *h*) or ‘numbers’ (*A* > *h*). Therefore, the contribution of predation-based mortality density-dependence (*DD*_*PM*_ = ∂[-*c*(*A*)*S*]/∂*A*) is also hump-shaped, intersecting zero at *h*.

Along gradients of *K* and *d*_*S*_, positive *DD*_*A*_ stems from each parameter’s direct and/or indirect influence on self-limitation (*DD*_*SL*_) and predation mortality (*DD*_*PM*_) components, mediated by densities. Equilibrial densities of resources and hosts are:

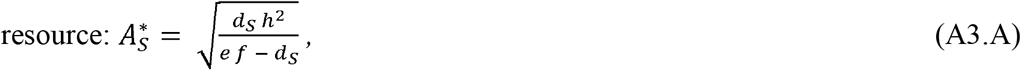

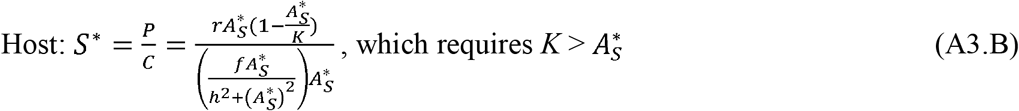

where *A*_*S*_* (eq. A3.A) sits at the host’s minimal resource requirement, unchanging with *K* (Fig. 2C) but increasing with *d*_*S*_ (Fig. A2C). Host density (*S*\*) is the ratio of primary production (*P* = *r A*_*S*_* [1 - *A*_*S*_*/*K*]) to per host resource consumption, *C* = *c*(*A*_*S*_*) *A*_*S*_* (eqs. A1.A, A3.B). Host invasion and feasibility of the *AS* equilibrium (i.e., positive densities of both species) requires that resources at the ungrazed boundary (*A*_*b*_* = *K*) exceed the host’s minimal resource requirement (i.e., *K* = *A*_*S*_* is the transcritical bifurcation *S*_*in*_). Host density, *S*\*, increases but saturates with increasing *K* since *P* increases with *K* (Fig. 2C). In contrast, *S*\* changes non-monotonically with *d*_*S*_, first decreasing, then increasing, to then finally decreasing again to zero as both *P* and *C* change shape with *A*_*S*_* (Fig. A2C).

#### Oscillations with host death rate in the disease-free case (Fig. A2)

Oscillations commence when equilibrium host density (*S*^*\**^) increases (rather than decreases) with its background death rate (*d*_*S*_; Fig. A2D). This phenomenon is best explained using relative sensitivities of production (*P*) and consumption (*C*) to a change in *A*, given that: (1) equilibrium host density is the ratio of resource production to consumption (*S*\* = *P*/*C*; eq. A3.B); that (2) growth rate of the resource is production minus total consumption (*G*_*A*_ = *P* – *C S*; after eq. A1.A); that (3) the resource’s self-feedback (*J*_*AA*_) is defined as the sensitivity of its growth rate to a change in its density (*J*_*AA*_ = ∂*G*_*A*_/∂*A*); and that (4) the resource at equilibrium (*A*_*S*_*) increases with death rate of the host (∂*A*_*S*_*/∂*d*_*S*_ > 0 always; eq. A3.A). With these four premises, parallel expressions describe change in host density (*S*\*) and self-regulation (*J*_*AA*_) with death rate (*d*_*S*_):

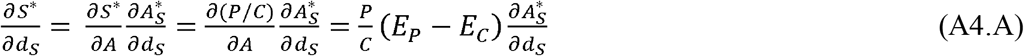

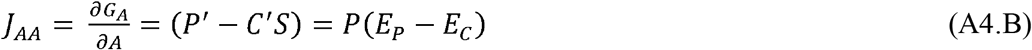

where *X*′ is the sensitivity of *X*\* to *A* (*X*′ = ∂*X*\*/∂*A* and *E*_*X*_ = *X*′/*X* is a relative (per unit) sensitivity. With those definitions, the host increases with its death rate (∂*S*\*/∂*d*_*S*_ > 0) when the relative increase in production, *E*_*P*_, due to higher resource density (stimulated by a release of foraging pressure from increased *d*_*S*_) exceeds the relative increase in consumption needs, *E*_*C*_, (i.e., when *E*_*P*_ > *E*_*C*_; eq. A4.A; yellow region, Fig. A2D). When this happens, the resource experiences self-facilitation (*J*_*AA*_ > 0 hence *DD*_*A*_ > 0; eq. A6.B; yellow region, Fig. A2E) and the *AS* system oscillates. Conversely, when the host declines with higher death rate (∂*S*\*/∂*d*_*S*_ < 0), the resource experiences net self-limitation (*J*_*AA*_ < 0 hence *DD*_*A*_ < 0) and the system is stable (blue regions, Fig. A2C,E). With high enough *d*_*S*_, however, the host can no longer persist (as now *A*_*S*_^*\**^ > *K*; white, Fig. A2C,E).

#### Insights into the epidemic equilibrium (and more on Fig A2)

Feedback loops are all evaluated at the equilibrium densities for each compartment (*A*^*\**^, *S*^*\**^, *I*^*\**^, *Z*^*\**^) at the equilibrium in question. In epidemic cases, we also defined total hosts as *H*^*\**^ = *S*^*\**^ + *I*^*\**^. Total host density (*H*^*\**^) can be determined by updated resource production (*P*) and per host resource consumption (*C*) terms that account for the epidemic compartments:

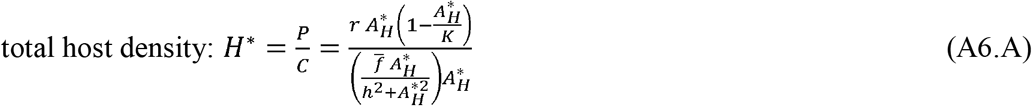

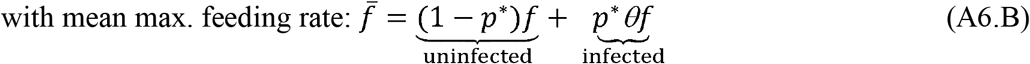

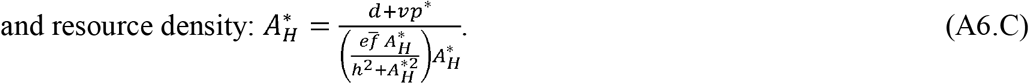

Here, maximum feeding rate 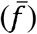 is updated as a prevalence-weighted average, where prevalence of infection (*p*\*) is the ratio of infected to total hosts (*p*\* = *I*\* / *H*\*). As for the disease-free case, *H*\* increases with *d*_*S*_ (Fig. A2D) when the relative sensitivity of *P* to *A* (*E*_*P*_) exceeds that of prevalence-weighted consumption (*E*_*C*_; so ∂*H*\*/∂*d*_*S*_ > 0 when *E*_*P*_ > *E*_*C*_). In this case, the resource again experiences positive density-dependence (*DD*_*A*_ > 0 so *J*_*AA*_ > 0; Fig. A2.F). When hosts decline with *d*_*S*_, (when *E*_*P*_ < *E*_*C*_) resources experience negative density-dependence (*DD*_*A*_ < 0 [Fig. A2] so *J*_*AA*_ < 0). Hence, in the region with oscillations (yellow, figs. A2D, F), the resource experiences positive density-dependence from the *DD*_*PM*_ term.

#### More about invasion of the parasite into a cycling AS system (Fig. A3)

We double-checked that the parasite could still invade the cycling resource-host region for variant 1. The disease subsystem (*I* and *Z*) invades if the net reproductive ratio *R*_0_ > 1. This net reproductive ratio – the ratio of propagule gains from new infections following release to losses from propagule death or consumption – was found using the next generation matrix [NGM] method (Diekmann et al. 2010), where:

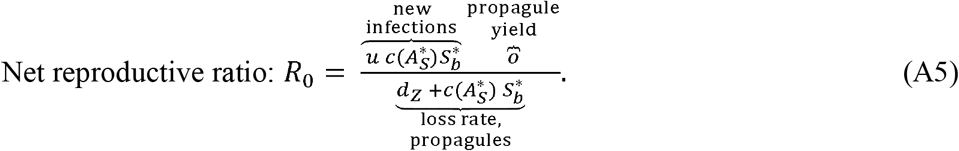

In regions where the *AS* disease-free system oscillates, net reproductive ratio must be calculated across the entire *AS* cycle (Bate and Hilker 2013; hence using mean host density 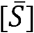 and resource density [*Ā*] (i.e., time averaged over a cycle period [of length τ] rather than at the *AS*-system’s boundary equilibrium, *S*_*b*_*; see Fig. A3). That invasion criterion is shown as a red line (Fig. A3). However, when the parasite invades a cycling *AS* system, *R*_0_ must exceed one averaged over an *AS* cycle (so, 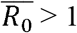 is required). We calculated mean *R*_0_ over 5,000 time steps (after first simulating 2,000 time steps to remove any possible transient behavior). All regions of black denote 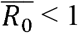 (Fig. A3). With finer gridding, we would find that 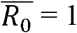 when *R*_0_ = 1 calculated at the equilibrium point. The germane point here, however, is that within the cycling *AS* zone (within the dotted *AS* Hopf), 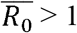. Hence, the cycling *AS* system posed no additional barrier to parasite invasion.

### Section 3: More on loop tracing and feedback at levels 1-4 (*F*_1_ – *F*_4_)

This part of the appendix provides more explanation of the levels of feedback used in loop tracing (Fig. 1). The analysis of feedback loops starts with a matrix of direct effects derived from the growth rate equations for each species (eq. 1; depicted in Fig. 1A). This matrix, the Jacobian (**J**), is composed of elements (*J*_*XY*_) linking one ‘species’ (i.e., species, class of host, stage of parasite, etc.) to another. Specifically, each element is the direct effect of a tiny increase in the density of one (*Y*) on the population growth rate of another (*G*_*X*_), so *J*_*XY*_ = ∂*G*_*X*_ / ∂*Y*, and where *J*_*XX*_ is the intraspecific, self-regulatory effect of a species on its own growth rate. The diagram for the disease system (*ASIZ*; Fig. 1A) composed of resources (*A*), susceptible hosts (*S*), infected hosts (*I*), and parasite propagules (*Z*) can be converted into a corresponding Jacobian (**J**_**ASIZ**_), in order of *A, S, I*, and *Z*, shown here for variant 3 (type III clearance rate and resource-dependent propagule yield, i.e., with maximal numbers of direct effects connecting species):

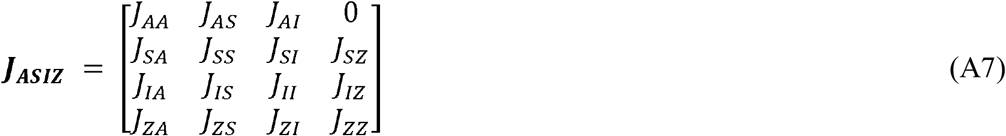

where each row describes the direct effect of the species in each column on population growth rate of the species in each row. Such a four species matrix has four corresponding levels of feedback (Fig. 1B-E). Feedback tracks the net effect of an increase of a species on its own density. Since that feedback can be looped through one, two, three, or four species, there are four levels of feedback. The first (*F*_1_) involves the sum of each species’ direct, self-regulatory effects on their own growth rate, where *J*_*XX*_ < 0 would indicate self-inhibition and *J*_*XX*_ > 0 indicates self-facilitation (Fig. 1B). Since feedback at level one (*F*_1_) sums these direct, intraspecific effects, it is the trace (Tr) of the Jacobian:

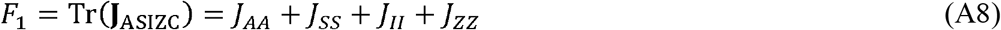

Level two feedback (*F*_2_) sums feedback within binary pairs of each species (Fig. 1C). For instance, feedback between the resources (*A*) and the susceptible host (*S*) would be the negative determinant of the *AS* subsystem (**J**_**AS**_), i.e., the Jacobian elements containing only *A* and *S*:

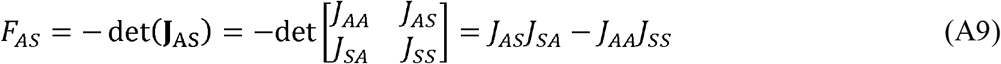

which is the host-resource interspecific interaction (*J*_*AS*_ *J*_*SA*_) minus the joint self-regulatory terms (*J*_*AA*_ *J*_*SS*_). Total level two feedback (*F*_2_) for the disease model (*ASIZ*) sums six such binary terms, three resource-dependent (*F*_*AS*_ + *F*_*AI*_ + *F*_*AZ*_) and three epidemiological (*F*_*SI*_ + *F*_*IZ*_ + *F*_*SZ*_):

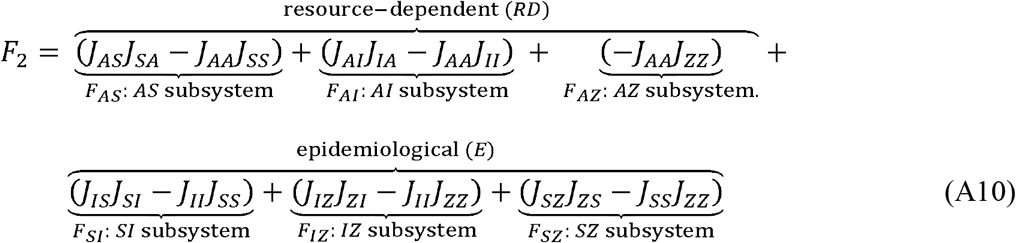

where *S* is the basal resource of the epidemiological loops. As pointed out in the main text, *RD* loops contain self-regulation of the resource (*J*_*AA*_) in each binary subsystem; the *E* loops do not. Level three feedback (*F*_3_) builds on these binary loops, combining interactions between trios of species and adding a new structure, the three species ‘net effect’ (Fig. 1D). Such a net effect has the structure *N*_*XY*.*Z*_ = *J*_*XY*_ (-*J*_*ZZ*_) + *J*_*ZY*_ *J*_*XZ*_, which reads as the net effect of *Y* on *X* via *Z*. It sums the direct effect of *Y* on *X, J*_*XY*_ (-*J*_*ZZ*_), and the indirect one via *Z, J*_*ZY*_ *J*_*XZ*_. Hence, *N*_*AS*.*I*_ is the net effect of *S* on *A* via *I*, that is the direct effect, *J*_*AS*_ (-*J*_*II*_), plus the indirect one, *J*_*IS*_ *J*_*AI*_. With this building block, the structure of feedback in, say, the resource-susceptible host-infected host (*ASI*) subsystem, is the determinant of a Jacobian just for this subsystem, **J**_**ASI**_:

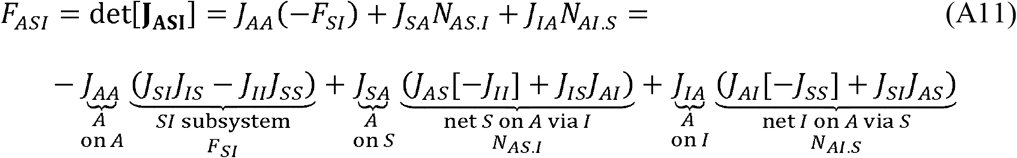

as written from the perspective of feedback on the basal, ecological resource (*A*) following increase in its density. The first term is intraspecific, with the direct effect of *A* on itself (*J*_*AA*_) times feedback in the non-interacting species (-*F*_*SI*_). The second is the interspecific direct effect of *A* on *S* (*J*_*AS*_) times the net effect of *S* back on *A* via *I* (*N*_*AS*.*I*_). The third is the interspecific direct effect of *A* on *I* (*J*_*AI*_) times the net effect of *I* back on *A* via *S* (*N*_*AI*.*S*_). Repeating this pattern, level three feedback (*F*_3_) sums the feedback in four trios:

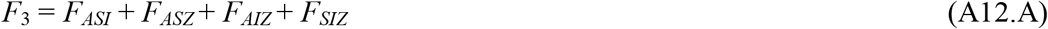

with:

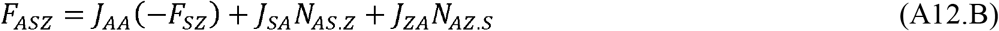

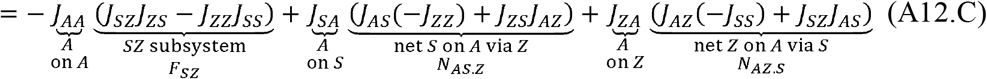

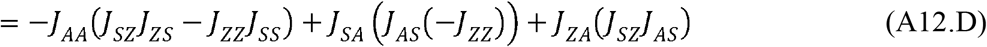

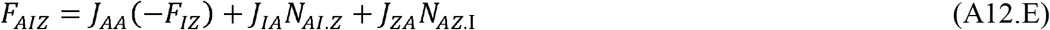

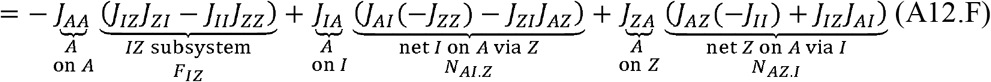

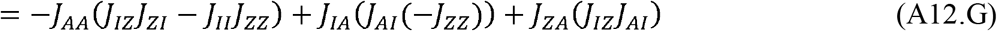

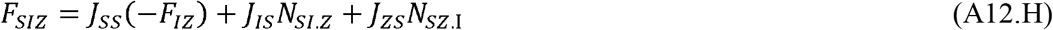

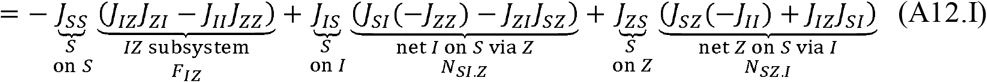

where *F*_3_ sums feedback in the resource-host (*ASI*), resource-susceptible-propagule (*ASZ*), resource-infected-propagule (*AIZ*), and disease-only (*SIZ*) subsystems (Fig. 1D). Note that some terms simplify (e.g., eq. A12.F to A12.G) because *J*_*AZ*_ = 0. Each three species net effect follows the *N*_*XY*.*Z*_ format above, and *F*_*SIZ*_ is derived from the perspective of the epidemiological resource, *S* (i.e., the feedback on *S* following a small increase in *S*).

Level four feedback (*F*_4_) or total system feedback sums the four species connections in the *ASIZ* disease system. It also determines if a disease equilibrium is potentially stable (negative feedback, *F*_4_ < 0) or a saddle (positive feedback, *F*_4_ > 0) in variant 2. Stability also depends on the oscillation criteria [*HC*], as in variant 1; *HC* pits *F*_1_ through *F*_3_ against *F*_4_ (eq. 7). It builds on the logic of the previous levels of feedback and is calculated from the negative determinant of **J**_**ASIZ**_; hence, *F*_4_ is:

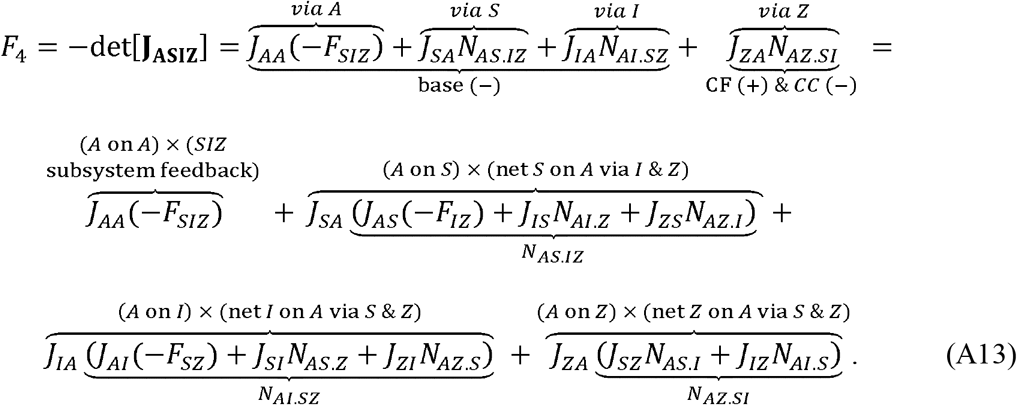

Level four feedback derived from the perspective of the basal resource (*A*), then, is the sum of the self-regulation of the resource (*J*_*AA*_) times feedback in the disease subsystem (-*F*_*SIZ*_) and the interspecific effect of *A* on *S, I*, and *Z* (*J*_*SA*_, *J*_*IA*_, *J*_*ZA*_) times their respective four species net effects back onto the resource (*via S, via I*, and *via Z* components, respectively; Fig. 1E). These four-species net effects follow a pattern resembling level three feedback. Take *N*_*AS*.*IZ*_, the net effect of *S* back on to *A* via *I* and *Z*. That term sums the direct interspecific effect of *S* on *A* (*J*_*AS*_, times feedback in the non-interacting *IZ* subsystem, -*F*_*IZ*_); the direct effect of *S* on *I* times the three species net effect of *I* back onto *A* via *Z, N*_*AI*.*Z*_; and the direct effect of *S* on *Z* times the net effect of *Z* back onto *A* via *I, N*_*AZ*.*I*_. In other words, in this part of *F*_4_, the resource affects *S*, and then the four species net effect traces the paths that *S* affects *A*, directly through consumption, or indirectly through the disease system (*I* and *Z*). Then, to complete *F*_4_: *A* directly affects *I*, then *I* net returns back to *A* directly or routed through *S* and *Z*; and *A* directly affects *Z*, and *Z* net returns back to *A* routed through *S* and *I* alone (since propagules have no direct effect on resources [*J*_*AZ*_ = 0] in all model variants). The loops underlying ‘base’ feedback and the cascade-fueling (*CF*) and consumption-constraint (*CC*) loops are also noted accordingly (Fig. 4).

As shown in the loop figure (Fig. 1), model variant 1 (with resource-dependent clearance rate, hence exposure rate) contains loops created by directly linking resources to the infected host (*J*_*IA*_) while variant 2 does not. Hence *F*_4_ for variant 1 contains *‘via I’* loops while that for variant 2 does not. Similarly, variant 2 (resource-dependent propagule yield) contains loops that contain direct links between resources and propagules (*J*_*ZA*_) while variant 1 does not. Hence, variant 2 contains *‘via Z’* loops its *F*_*4*_ while that for variant 1 does not. Variant 3 contains all of the loops above.

### Section 4: More on loop tracing of variants 1, 2, and 3

#### More about level three feedback in variant 1 (Fig. A4)

In variant 1, oscillations begin just about when the resource experiences positive density-dependence (*DD*_*A*_ > 0) hence self-facilitation (*J*_*AA*_ > 0; Fig. 2D, F). However, oscillations are not born from *J*_*AA*_ alone. Instead, *J*_*AA*_ influences feedback in the intraspecific part of the resource-susceptible host-propagule (*ASZ*) three species subsystem (Fig. 3C-E); then, *F*_*ASZ*_ shapes level three feedback (Fig. 3B); and then, *F*_3_ transitions from negative to positive just about where the Hopf arises (Fig. 3A). *J*_*AA*_ > 0 triggers oscillations through its influence on the *ASZ* subsystem but not the two others containing *J*_*AA*_ (Fig. A4A) because feedback in its *SZ* subsystem (*F*_*SZ*_ *= J*_*SZ*_ *J*_*ZS*_ - *J*_*SS*_ *J*_*ZZ*_) is so strong. That strength of *F*_*SZ*_ stems from strong self-inhibition of *Z* and especially strong self-inhibition of *S* (i.e., *J*_*ZZ*_ and *J*_*SS*_ have large magnitude, hence *J*_*SS*_ *J*_*ZZ*_ is large) in combination with weak interspecific effects (so *J*_*SZ*_ *J*_*ZS*_ is small; Fig. A2B). The other two subsystems containing *J*_*AA*_ (parts of *AIZ* and *ASI*) each have two species subsystems (*IZ* and *SI*, respectively) that provide little to no feedback; in both cases, infected hosts (*I*) experience very little self-inhibition (i.e., *J*_*II*_ is small), thereby rendering these two subsystems (*F*_*IZ*_ and *F*_*SI*_) and the three-species subsystems enveloping them largely irrelevant (Fig. A4C, D; in fact *F*_*SI*_ = 0).

#### More about level four feedback in variant 2

Resource-related net positive feedback in *F*_*4*_ lies at the heart of analysis of alternative stable states in model variant 2. The ‘base’ level feedback sums the ‘*via A*’ component (*J*_*AA*_ (− *F*_*SIZ*_)) the ‘*via S*’ component (*J*_*SA*_ *N*_*AS*.*IZ*_), as there is no ‘*via I*’ component (since *J*_*IA*_ = 0; Fig. 1.E, eq. A13). This sum of ‘*via A*’ and ‘*via S*’ into ‘base’ feedback is negative (Fig. 4). If propagule yield remains constant (hence *J*_*ZA*_ = 0), this base component of *F*_4_ provides negative feedback. However, variant 2 showcases implications of breaking this assumption (*J*_*ZA*_ > 0), which thereby introduces the ‘via Z’ (*J*_*ZA*_ *N*_*AZ*.*SI*_) loops to *F*_4_. We separate out these terms into the one generating positive and negative feedback (Fig. 1E, 4E-G):

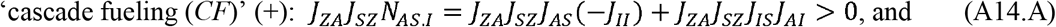

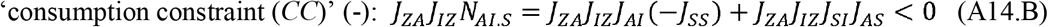

where the ‘cascade fueling’ (*CF*) component generates positive feedback because resources increase spores that depress susceptible host density epidemiologically (*J*_*SZ*_ < 0), thereby releasing resources (since the net effect of *S* on *A* via *I* is negative [*N*_*AS*.*I*_ < 0]; eq. A14.A). In other words, in *CF*, resources themselves fuel the parasite-driven trophic cascade that leads to more resource release. In contrast, the ‘consumption constraint’ (*CC*) component generates negative feedback because resources increase infected hosts (epidemiologically) that net consume resources (via *N*_*AI*.*S*_; eq. A14.B). At the saddle equilibrium, positive feedback arises because the *CF* component (+) exceeds the two other stabilizing ones (*CC* + basal).

#### ‘Loop tracing’ in variant 3 (Fig. A5)

In the variant with resource-dependent exposure and propagule yield, as the Hopf criterion changes sign, *F*_1_ and *F*_2_ both weaken strongly (Fig. A5A). Not surprisingly, within the intraspecific, self-regulatory *F*_1_ terms, *J*_*AA*_ shifts positive almost exactly as *HC* does (Fig. A5B). However, strongly stabilizing (large negative) effects of propagules (*J*_*ZZ*_) ensure *F*_1_ stays quite negative and stabilizing (Fig. A5B). Within *F*_2_, the ‘resource-dependent’ (*RD*) terms (involving *A*: *F*_*AS*_, *F*_*AI*_, *F*_*AZ*_) weaken with *d*_*S*_ while ‘epidemiological’ (*E*) ones (*F*_*SI*_, *F*_*IZ*_, *F*_*SZ*_) strengthen (become more negative; see also eq. A12). Focusing on those resource-dependent loops, we see that *F*_*AZ*_ weakens most similarly to *F*_2_ (Fig. A5C). Finally, within *F*_*AZ*_, *J*_*AA*_ weakens and changes sign while *J*_*ZZ*_ acts as a magnifier. So, consumption-driven density-dependence of resources (via *J*_*AA*_) flips positive almost as oscillations commence (with *HC* > 0). However, it operates by weakening *F*_1_ and *F*_2_ via its relationship to feedback with a resource-dependent binary loop involving propagules (in *F*_*AZ*_).

### Section 5: Complex system behaviors in variant 3

Type III clearance and resource-dependent propagule yield also introduce more exotic regimes beyond those in simpler variants (Fig. A6). For instance, at high carrying capacity of the resource (*K*) and high death rate of susceptible hosts (*d*_*S*_), a second region of alternative stable states (Alt-SS [2]) demonstrates epidemics at either stable equilibrium or a stable limit cycle (oscillations). To illustrate, a sufficiently large drop of the resource (to *A*_0_ = 0.5 mg/L at time *t* = 50 days) shifted stable disease dynamics to oscillatory ones (Fig. A6.B). However, stable and oscillating epidemic states are separated by an unstable cycle (rather than a saddle point as in Alt-SS[1]). Both the stable and unstable cycles were introduced by the limit point cycle bifurcation (like a fold, but for cycles; Fig. A6.A). Oscillations in Alt-SS (2) resemble those of variant 1 (Fig. 2B), but the propagule cycle has two peaks because its hinges on two resources. Like variant 1, peak ecological resource (*A*) leads to peak epidemiological resource (*S*) which fuels rise in both propagules (*Z*, to its maximum peak) then infected density (*I*). The path to peak *I* crashes *A*, then *S*. The second, smaller peak stems from subsequent rapid rise in *A* which increases yield of propagules from the few remaining infected hosts. This smaller *Z* peak cannot create another full epidemic wave; rather, it seeds the return of the epidemic as hosts rebound.

A third type of alternative states, Alt-SS (3), arises where Alt-SS (1) and epidemic oscillations collide. At slightly higher *d*_*S*_, this new region produces either stable host-resource (*AS*) or oscillatory disease (*ASIZ*) dynamics, depending on initial conditions. A large enough introduction of propagules to the stable *AS* system (*Z*_0_ = 100 #/L at *t* = 50 days, Fig. A6.C) can start an epidemic. This epidemic regime demonstrates even more complex dynamics than those previously demonstrated. The oscillating *ASIZ* attractor undergoes a period-doubling route to chaos with increased *K* (Fig. A6.D), eventually yielding a weakly chaotic attractor (Fig. A6.E). At lower *K*, the base oscillation largely follows that described for variant 1 (Fig. 2B). Higher *K* leads to a two-peak cycle, one that exacerbates the two-resource phenomenon of Alt-SS (2). Once again, high peak density of the basal resource (*A*) fuels high peak of the epidemiological resource (*S*), then high peak propagule density (*Z*) followed by high peak infected host (*I*). That sequence leads to large depletion of *A*. The recovery of *A* then fuels the small peak of propagules that seeds the small peak in infected hosts. However, the epidemic stays at the small peak because the epidemiological resource (*S*) is still declining, thereby undermining disease transmission. The small peak in *I* depresses *A* again, but full recovery of both resources (*A*, then *S*) coupled with seeding of Z by the small peak, pushes the full cycle to renew.

Hence, the two-peak cycle extends longer (i.e., it is ‘period-doubled’). It alternates a high peak (created when recovery of both resources fuel maximal transmission) vs. a small peak (created when partial recovery of the basal resource raises transmission, but still-declining host density depresses transmission, smothering full increase). Seeding events of this kind involving both ecological and epidemiological resources of the epidemic become more complicated and varied with further increases of *K*. Eventually, the cycles proceed entirely aperiodically. Without a regular pattern of the dynamics, it becomes more difficult to predict the future of the epidemic. The series of period-doubling bifurcations with increased enrichment is known as the period-doubling (intermittency) route to chaos, a statistically distributed series of period-doubling bifurcations where the period eventually approaches infinity and is therefore chaotic (Strogatz, 2024).

Sensitivity to initial conditions is the hallmark of chaotic attractors; a minute difference in initial conditions will yield wildly different behavior as the two trajectories spread differently across the chaotic attractor. This sensitivity is captured by measuring the first Lyapunov exponent *λ*, which describes the rate of separation of two infinitesimally close trajectories (Strogatz, 2024). That rate is exponential in the case of a chaotic attractor, so the difference between two trajectories *δ* over time is defined as:

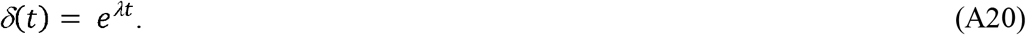

When plotted on a log scale, this separation appears as a positive linear slope where that slope defines the Lyapunov exponent *λ*. The slope asymptotes over time at the diameter of the chaotic attractor, which defines the maximum separation between two trajectories on that attractor. Both the slope defining *λ* and that asymptote are shown in the right panel of Fig. A6.E. The positive LCE slope measured here (∼ 0.005) indicates weak chaos. As *K* continues to increase, the reverse pattern occurs; subsequent period-doubling bifurcations collapse the many-peaked dynamics to simple, single-peaked oscillations (not shown). Plots in Fig. A6.E were generated with the help of software described in Sandri (1996).

In the main text, we applied loop analysis the first period doubling bifurcation on the route to chaos. To do so, we examined Floquet multipliers. Floquet multipliers are the periodic cousins of eigenvalues, which underlie all the stability analysis conducted to this point. Briefly, eigenvalues *λ* for a matrix of equations (such as the Jacobian matrix) are determined from the characteristic equation of the system. For Jacobian matrix **J**, the characteristic equation is given by solving the determinant of **J** – *λ I* = 0, where I is the identity matrix. For stability in systems of ordinary differential equations, such as our epidemic models, all eigenvalues must have negative real parts. Notably, the feedback loops we dissect in this manuscript are related to eigenvalues via the characteristic equation, where level 1 loops are unscaled, level 2 loops are scaled by *λ*, level 3 loops are scaled by *λ*^*2*^, and so on. We generally solve for eigenvalues analytically, but in cases that are analytically intractable, they can be solved for numerically.

Floquet multipliers determine stability on periodic solutions to differential equations. They are analogous to eigenvalues but can never be solved for analytically. To determine floquet multipliers numerically, we first solve the matrix equation:

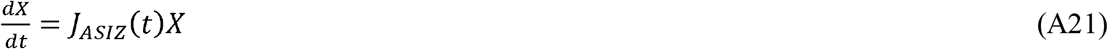

over one period (0 – *T*) with the identity matrix *I* as the initial condition (*X(0) = I*) and the states of each species initialized at some point on the periodic solution. Floquet multipliers are the eigenvalues to the solution to this equation *X(T)*, (the ‘monodromy’ or ‘fundamental matrix’; Grimshaw, 1993). Floquet multipliers, like eigenvalues, can be used to determine stability, albeit the criteria are slightly different. If all real parts of the Floquet multipliers are between -1 and 1, the orbit is stable. If any multipliers fall outside that range, the cycle is unstable. As such, period doubling of a cycle can be detected by examining when the dominant Floquet multiplier drops below negative one (whereas rising above one denotes a Neimark-Sacker bifurcation). In our variant 3, the dominant Floquet multiplier falls below negative one ∼*K* = 3.041 (Fig. 5C).

Because Floquet multipliers are related to eigenvalues, we can relate them numerically to feedback loops. Solving for eigenvalues of the fundamental matrix (*X(T)*, that is, the solution to eq. A21 after period *T* time) gives the Floquet multipliers of the orbit. Similarly, the fundamental matrix *X(T)* can be decomposed into its constituent loops to determine which of those loops drives the period doubling bifurcation, as we did in the main text. To do so, we computed each element of the monodromy matrix while solving equation A21. Future work must more rigorously determine the relationships between bifurcations on orbits and loop analysis given the importance of these bifurcations in ecological theory.

## APPENDIX FIGURES

**Figure A1:**
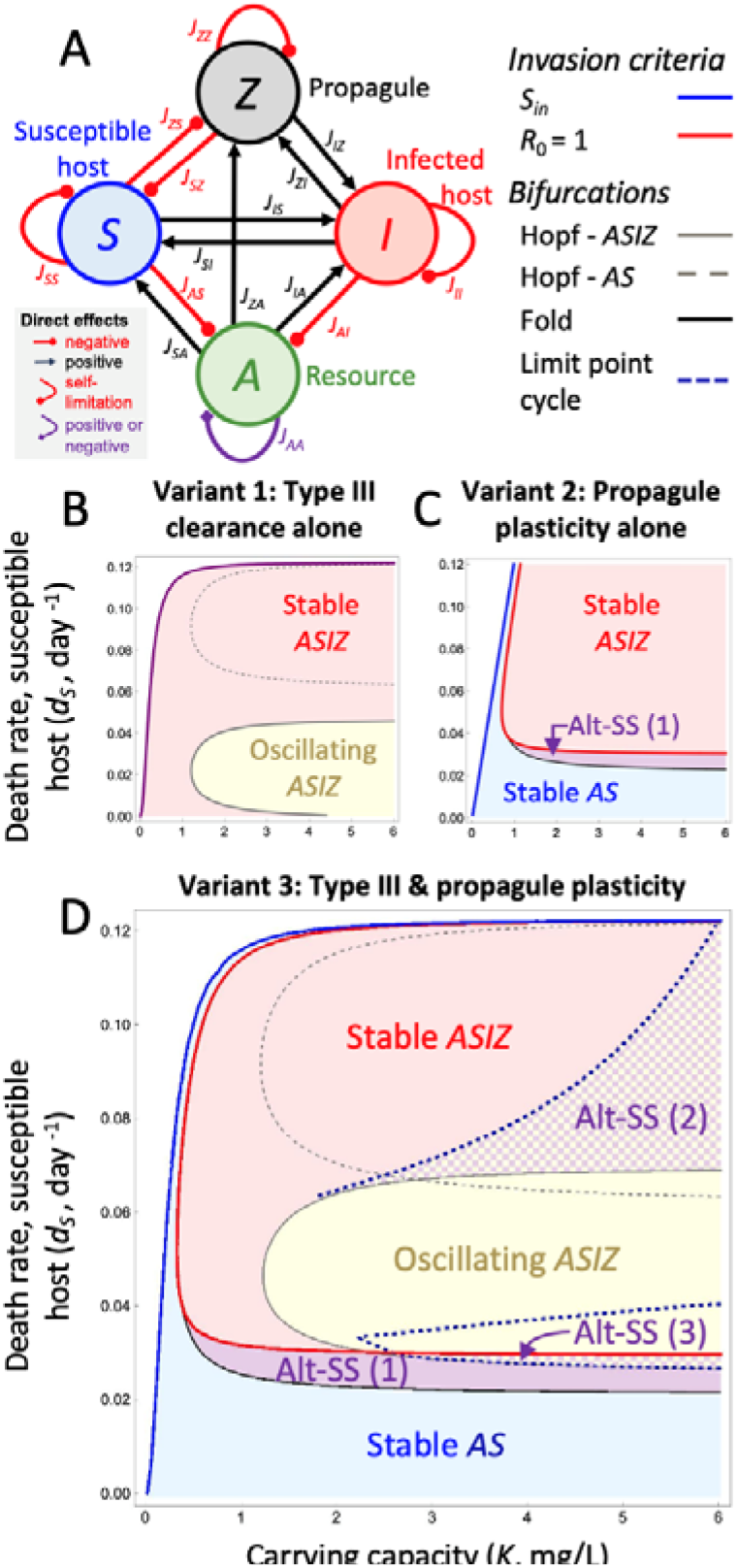
Dynamical regimes of three model variants of the model of resources, hosts, and parasites (*ASIZ*). Bifurcation maps along gradients of resource carrying capacity (*K*) and susceptible host death rate (*d*_*S*_). Hosts and parasites can invade past blue (‘*S*_*in*_’) and red (‘*R*_0_ = 1’) lines, respectively. Combinations of *K* and *d*_*S*_ produce stable host-only (*AS*, blue) and disease (*ASIZ*, red) dynamics, oscillations (yellow), and/or alternative states (purple). **(A)** Direct effects of the full disease (*ASIZ*) system. Black arrows indicate positive direct effects while red circles indicate negative ones, labeled with its corresponding Jacobian element (see text). **(B)** Variant 1: Type III clearance yields oscillations in the host-only (past ‘Hopf-*AS*’) and the disease system (past ‘Hopf-*ASIZ*’; yellow region). **(C)** Variant 2: Resource-dependence of propagule yield generates alternative stable states (purple; ‘Alt-SS (1)’), generated by a fold (black curve) between stable host-only (blue, *AS*) and disease (red, *ASIZ*) regimes. **(D)** Variant 3: Combining resource-dependent clearance and yield creates oscillations, bistability, and more complex behaviors. At higher *d*_*S*_, epidemics reach either stable or oscillatory dynamics in ‘Alt-SS (2)’. In ‘Alt-SS (3)’, a stable host-only regime exists with an oscillating disease regime (past the limit point cycle bifurcation [blue dashed line]: purple and yellow checkered). In here, a period doubling route to chaos can arise. (See also Table 1).

**Figure A2:**
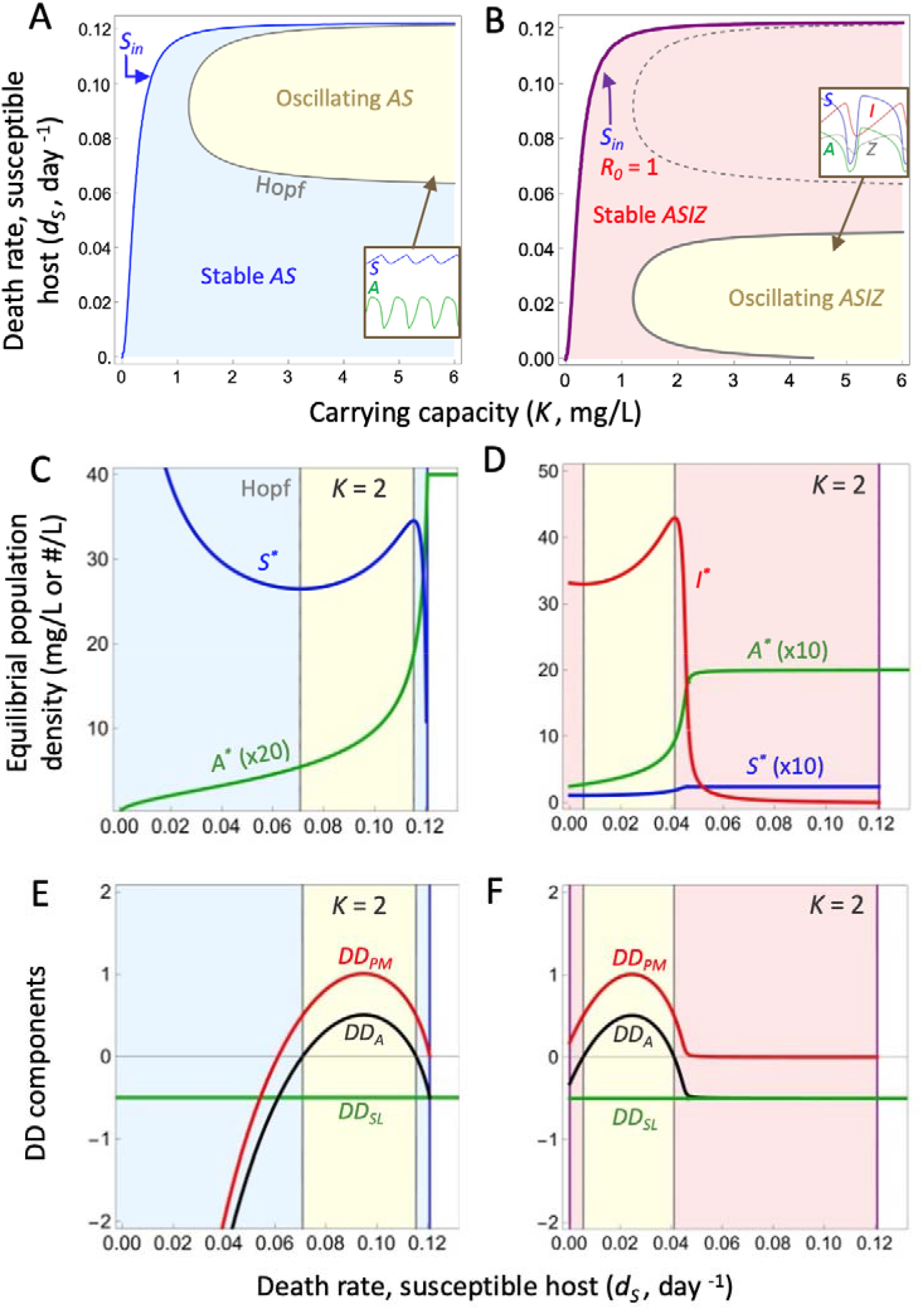
Positive density-dependence and oscillations with increasing host death rate *d*_*S*_. **(A)** and **(B)** are the same as Fig. 2. Densities of resources (*A*\*) and susceptible hosts (S*) are plotted with increasing *d*_*S*_ for the **(C)** non-epidemic and **(D)** epidemic system. Equilibria are plotted in the oscillation region, not cycle averages, as feedback is calculated on equilibria. Net density-dependence of the resource (*DD*_*A*_; black line) and its components of predation mortality (red: *DD*_*PM*_) and self-limitation (green: *DD*_*SL*_) can be plotted along gradients of *d*_*S*_ for the **(E)** non-epidemic and **(F)** epidemic system. When *DD*_*A*_ > 0, both systems oscillate. (non-epidemic *K* = 2, epidemic *K* = 2; see also Table 1).

**Figure A3:**
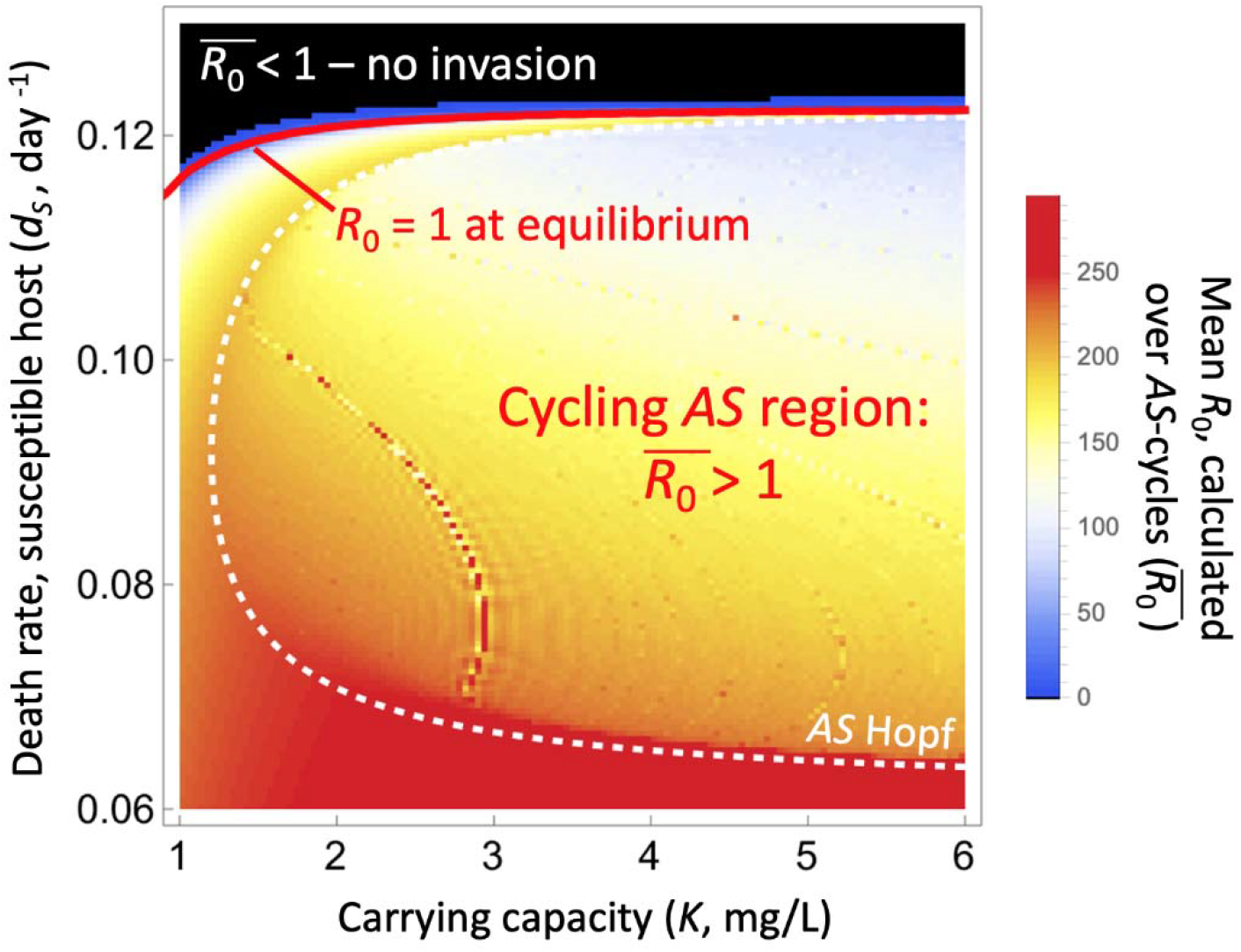
Calculation of mean net reproductive ratio on AS cycles in variant 1. Net reproductive ratio must be calculated across consumer-resource cycles after the *AS* Hopf (white dashed line) to determine the invasion of parasites, rather than relying on equilibrial densities (*R*_0_ = 1, solid red line). For each combination of carrying capacity of resource (*K*) and death rate of susceptible hosts (*d*_*S*_), we simulated the *AS* subsystem for a long period of time after it reached its stable limit cycle. *R*_0_ was calculated along the dynamics and averaged to determine (here blue to red colors indicate ≥ 1 while black indicates < 1). In this system, there was no change in invasion success (when rare) of the parasite into a cycling host and resource (within the *AS* Hopf; minor differences between edge of the black region and red *R*_0_ = 1 line involves gridding of the simulation).

**Figure A4:**
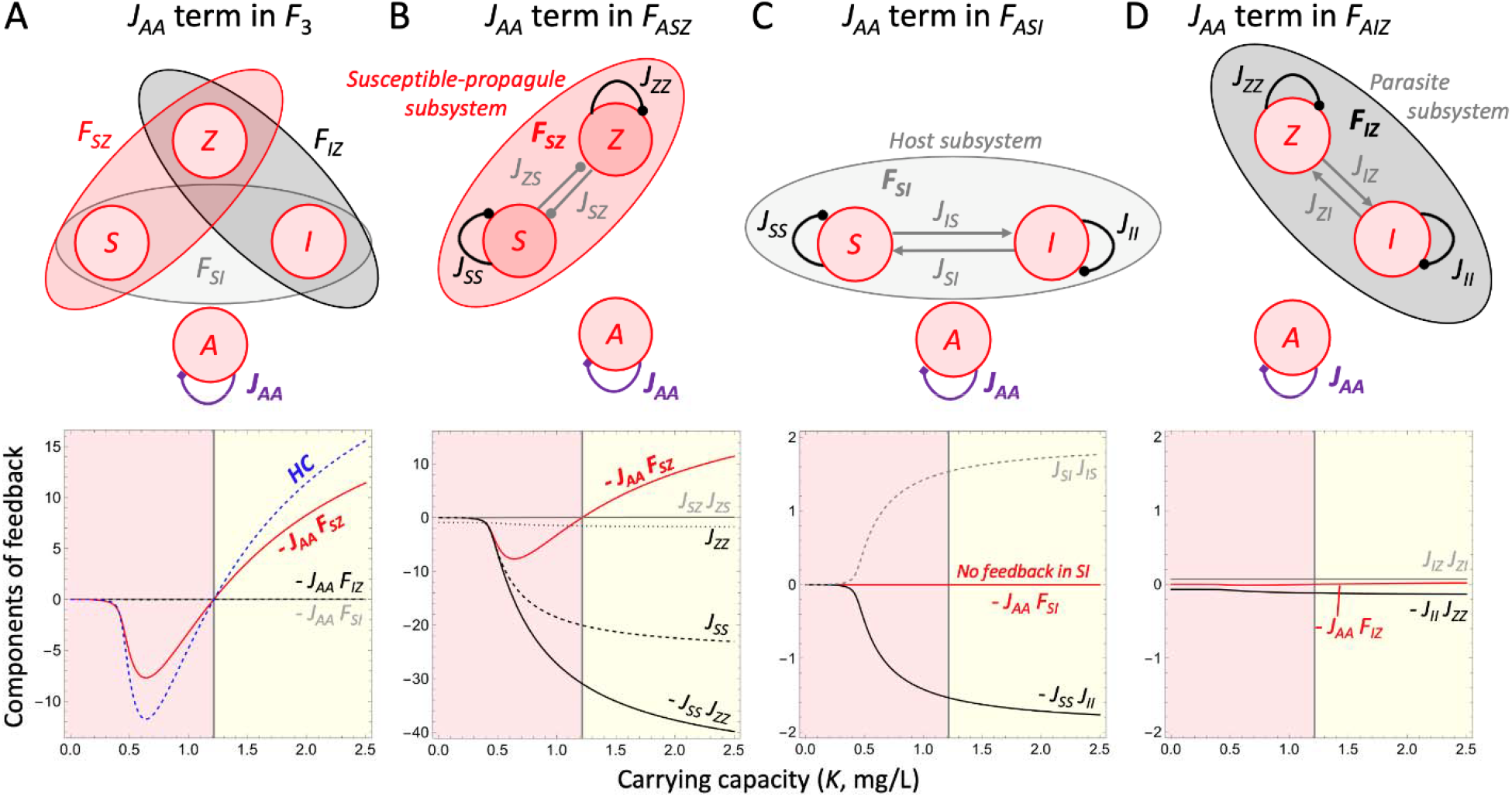
The influence of the ASZ subsystem in variant 1 oscillations. As shown in the main text (Fig. 4), the resource-susceptible-propagule (*ASZ*) subsystem contains **(A)** self-regulation of the resource (as -*J*_*AA*_ *F*_*SZ*_) and most strongly weakens level three feedback (*F*_3_), hence the Hopf criterion (*HC*) for oscillations. Two other subsystems, *ASI* and *AIZ*, also contain the *J*_*AA*_ term (as -*J*_*AA*_ *F*_*SI*_ and -*J*_*AA*_ *F*_*IZ*_, respectively), but they provide little feedback, hence exert little influence on *F*_3_. The difference lies in the non-interacting subsystems. **(B)** The *ASZ* subsystem is influential because it is proportional to very strong self-regulation by the susceptible host (*J*_*SS*_) and that of propagules (*J*_*ZZ*_). Combined, they yield strong joint intraspecific feedback; meanwhile, interspecific feedback (from *J*_*SZ*_ *J*_*ZS*_) is small. **(C)** The relevant *J*_*AA*_-containing component of the *ASI* subsystem, proportional to *F*_*SI*_, does contain *J*_*SS*_, but tiny feedback from infected hosts [*J*_*II*_ is close to zero]) lessens it while the inter-class component (*J*_*SI*_ *J*_*IS*_) counters it completely. **(D)** The component of *AIZ* subsystem with *J*_*AA*_ that is proportional to *F*_*IZ*_ is small because both joint intra- (*J*_*II*_ *J*_*ZZ*_) and inter-stage feedback (*J*_*IZ*_ *J*_*ZI*_) are small. (See also Table 1).

**Figure A5:**
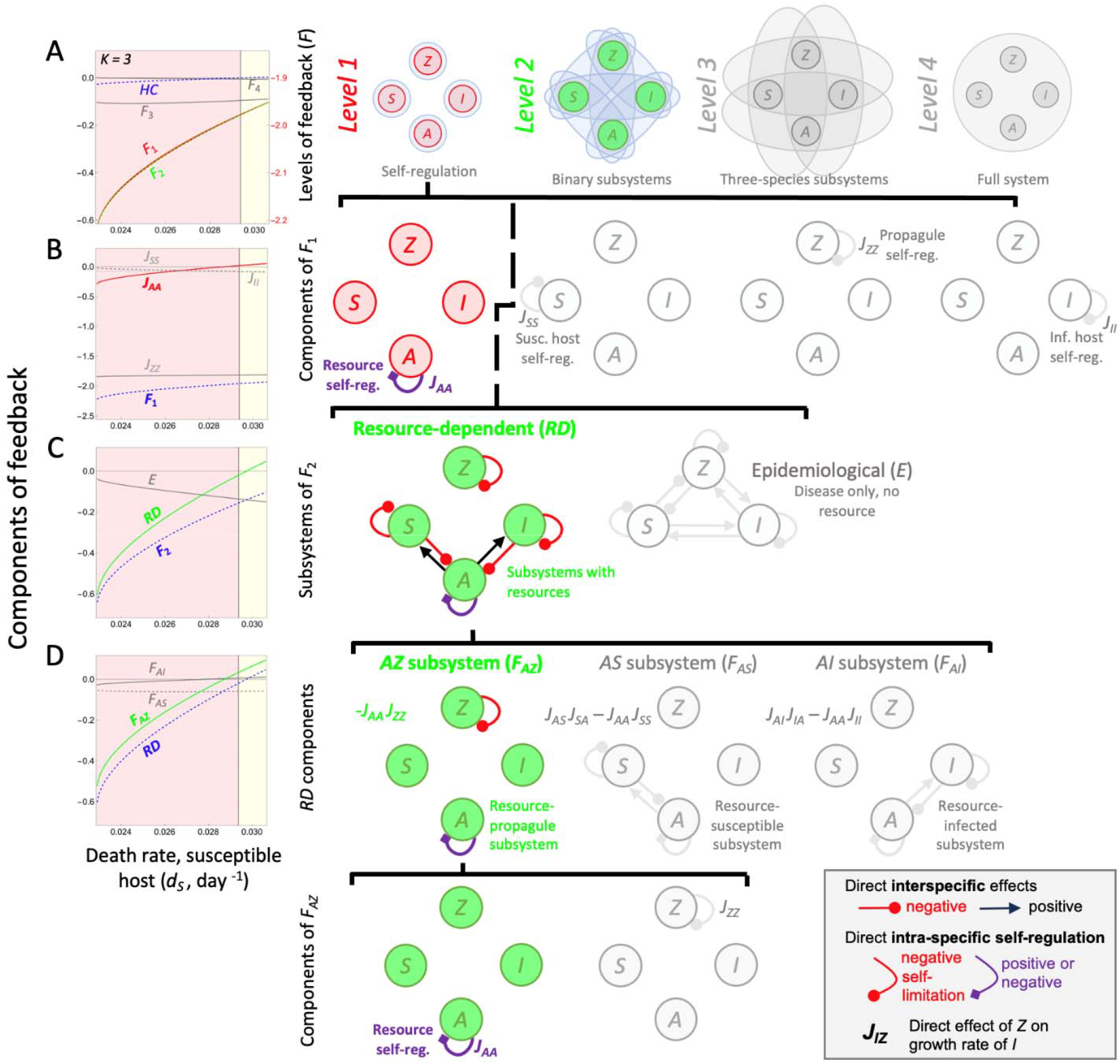
‘Loop tracing’ the genesis of oscillations in variant 3. Oscillations occur in the disease model (*ASIZ*) with type III clearance and resource-dependent propagule yield through the influence of density-dependence of the resource (*DD*_*A*_, hence *J*_*AA*_) on level 1 and 2 feedback. **(A)** Along a gradient of death rate of susceptible hosts (*d*_*S*_), the Hopf criterion (*HC*) flips positive as both *F*_1_ and *F*_2_ increase. Note that *F*_1_ follows scaling on the right y-axis. **(B)** Within *F*_1_, *J*_*AA*_ flips positive almost when *HC* does, but strong stabilization from spores (*J*_*ZZ*_) keeps *F*_1_ quite negative. (**C)** Within *F*_2_, a set of resource-dependent binary loops becomes weaker with *F*_2_, while a set of epidemiological ones strengthens. **(D)** Within those resource-dependent loops, resource-spore feedback (*F*_*AZ*_) exerts the most influence. Finally, within (*F*_*AZ*_), *J*_*AA*_ switches sign while *J*_*ZZ*_ provides magnitude, much like *J*_*ZZ*_ did (Fig. 3E). (See also Table 1).

**Figure A6:**
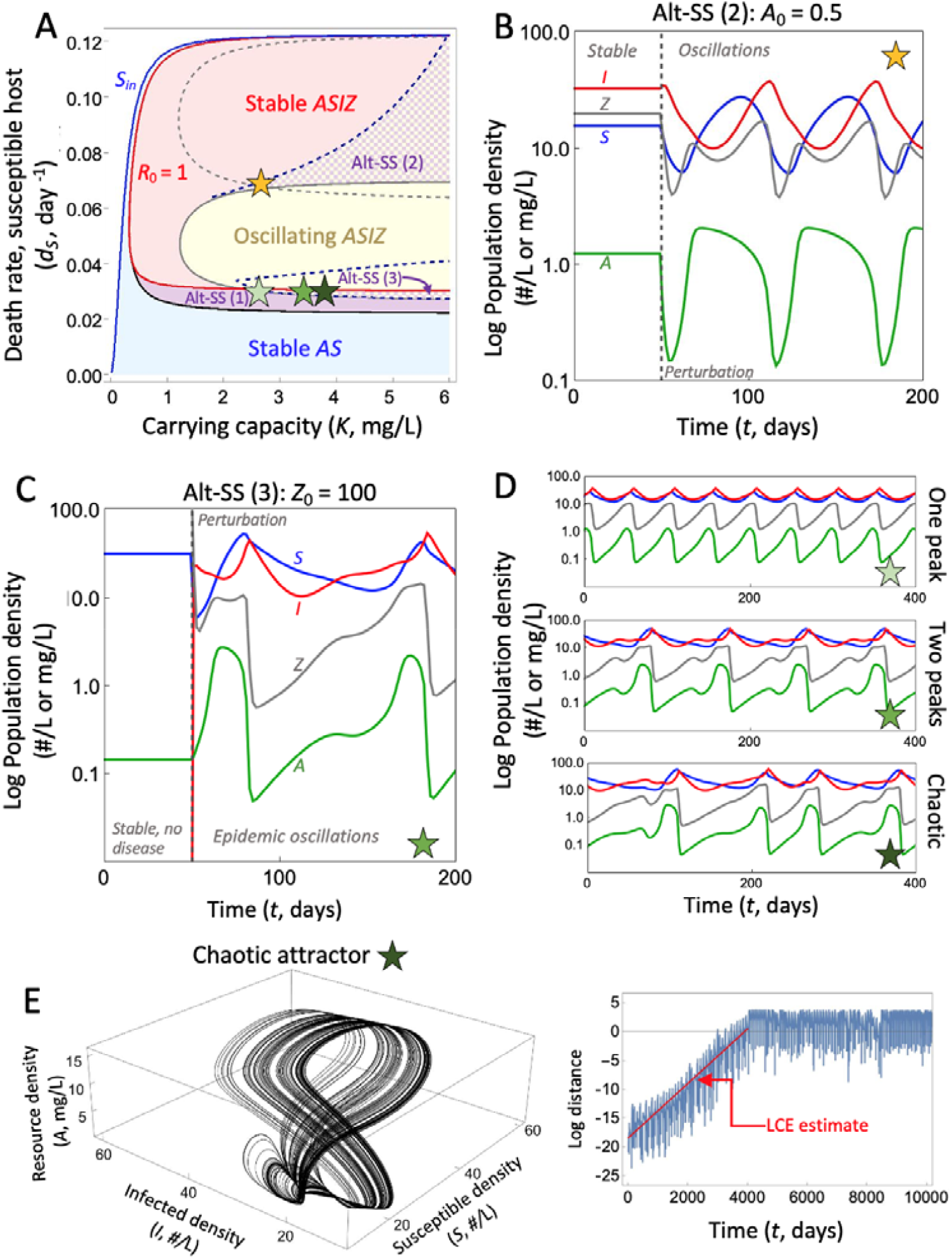
Complex behaviors in variant 3. **(A)** is the same as Fig. 5A. Bifurcation curves are denoted according to Fig. A1. Stars indicate various complex behaviors demonstrated in the following panels. **(B)** A new region of alternative stable states (Alt-SS [2]) occurs at high *K* and *d*_*S*_, separating stable and oscillating epidemic regimes. Time series demonstrate a shift between these regimes when the resource *A* was perturbed (*A*_*0*_ = 0.5). **(C)** A second new region of alternative stable states (Alt-SS [3]) occurs when oscillations collide with the original bistable region (Alt-SS [1]). Time series demonstrate a shift between a stable disease-free regime and an oscillating epidemic regime (here, initial perturbation of *Z*_*0*_ = 100). **(D)** The oscillating epidemic regime in Alt-SS (3) undergoes a series of period doubling bifurcations with increasing *K*, eventually yielding aperiodic, chaotic behavior. **(E)** We proved the existence of a weakly chaotic attractor by examining the slope of the first characteristic Lyapunov exponent (see Appendix section 5).

## REFERENCES

Althouse, Benjamin M., and Laurent Hébert-Dufresne. 2014. “Epidemic Cycles Driven by Host Behaviour.” Journal of The Royal Society Interface 11 (99): 20140575. 10.1098/rsif.2014.0575.

Anderson, Roy M., and Robert M. May. 1978. “Regulation and Stability of Host-Parasite Population Interactions: I. Regulatory Processes.” The Journal of Animal Ecology 47 (1): 219. 10.2307/3933.

Aron, Joan L., and Ira B. Schwartz. 1984. “Seasonality and Period-Doubling Bifurcations in an Epidemic Model.” Journal of Theoretical Biology 110 (4): 665–79. 10.1016/s0022-5193(84)80150-2.

Auld, Stuart K. J. R., Spencer R. Hall, Jessica Housley Ochs, Mathew Sebastian, and Meghan A. Duffy. 2014. “Predators and Patterns of Within-Host Growth Can Mediate Both Among-Host Competition and Evolution of Transmission Potential of Parasites.” The American Naturalist 184 (S1): S77–90. 10.1086/676927.

Bate, Andrew M., and Frank M. Hilker. 2013. “Complex Dynamics in an Eco-Epidemiological Model.” Bulletin of Mathematical Biology 75 (11): 2059–78. 10.1007/s11538-013-9880-z.

Bedhomme, Stephanie, Philip Agnew, Christine Sidobre, and Yannis Michalakis. 2004. “Virulence Reaction Norms across a Food Gradient.” Proceedings of the Royal Society of London. Series B: Biological Sciences 271 (1540): 739–44. 10.1098/rspb.2003.2657.

Blower, Sally, and Jonathan Roughgarden. 1987. “Population Dynamics and Parasitic Castration: A Mathematical Model.” The American Naturalist 129 (5): 730–54. 10.1086/284669.

Borer, Elizabeth T., Amy E. Kendig, and Robert D. Holt. 2023. “Feeding the Fever: Complex Host□pathogen Dynamics along Continuous Resource Gradients.” Ecology and Evolution 13 (7). 10.1002/ece3.10315.

Buck, Julia C., and William J. Ripple. 2017. “Infectious Agents Trigger Trophic Cascades.” Trends in Ecology & Evolution 32 (9): 681–94. 10.1016/j.tree.2017.06.009.

Cáceres, C.E., G. Davis, S. Duple, et al. 2014. “Complex Daphnia Interactions with Parasites and Competitors.” Mathematical Biosciences 258 (December): 148–61. 10.1016/j.mbs.2014.10.002.

Case, T. J. (2000). An illustrated guide to theoretical ecology (Nachdr.). Oxford Univ. Press.

Civitello, David J., Rachel M. Penczykowski, Jessica L. Hite, Meghan A. Duffy, and Spencer R. Hall. 2013. “Potassium Stimulates Fungal Epidemics in Daphnia by Increasing Host and Parasite Reproduction.” Ecology 94 (2): 380–88. 10.1890/12-0883.1.

Civitello, David J., Rachel M. Penczykowski, Aimee N. Smith, Marta S. Shocket, Meghan A. Duffy, and Spencer R. Hall. 2015. “Resources, Key Traits and the Size of Fungal Epidemics in DAphnia Populations.” Journal of Animal Ecology 84 (4): 1010–17. 10.1111/1365-2656.12363.

Cornet, Stéphane, Coraline Bichet, Stephen Larcombe, Bruno Faivre, and Gabriele Sorci. 2014a. “Impact of Host Nutritional Status on Infection Dynamics and Parasite Virulence in a Bird□malaria System.” Journal of Animal Ecology 83 (1): 256–65. 10.1111/1365-2656.12113.

Cornet, Stéphane, Coraline Bichet, Stephen Larcombe, Bruno Faivre, and Gabriele Sorci. 2014b. “Impact of Host Nutritional Status on Infection Dynamics and Parasite Virulence in a Bird□malaria System.” Journal of Animal Ecology 83 (1): 256–65. 10.1111/1365-2656.12113.

Cortez, Michael H. 2024. “Comparing the Differing Effects of Host Species Richness on Metrics of Disease.” Ecological Monographs 94 (4). 10.1002/ecm.1626.

Cotter, Sheena C, and Ekhlas Al Shareefi. 2022. “Nutritional Ecology, Infection and Immune Defence — Exploring the Mechanisms.” Current Opinion in Insect Science 50 (April): 100862. 10.1016/j.cois.2021.12.002.

Cressler, Clayton E., William A. Nelson, Troy Day, and Edward McCauley. 2014. “Disentangling the Interaction among Host Resources, the Immune System and Pathogens.” Ecology Letters 17 (3): 284–93. 10.1111/ele.12229.

Dambacher, J. M., Li, H. W., & Rossignol, P. A. (2003). Qualitative predictions in model ecosystems. Ecological Modelling, 161(1-2), 79–93. 10.1016/S0304-3800(02)00295-8

Diekmann, O., J. A. P. Heesterbeek, and M. G. Roberts. 2010. “The Construction of Next-Generation Matrices for Compartmental Epidemic Models.” Journal of The Royal Society Interface 7 (47): 873–85. 10.1098/rsif.2009.0386.

Fearon, Michelle L., Kristel F. Sánchez, Syuan□Jyun Sun, et al. 2025. “Resource Quality Differentially Impacts Daphnia Interactions with Two Parasites.” Ecosphere 16 (3). 10.1002/ecs2.70234.

Fussmann, Gregor F., Stephen P. Ellner, Kyle W. Shertzer, and Nelson G. Hairston Jr. 2000. “Crossing the Hopf Bifurcation in a Live Predator-Prey System.” Science 290 (5495): 1358–60. 10.1126/science.290.5495.1358.

Grover, J. P. (1997). Resource competition (Vol. 19). Springer Science & Business Media.

Hall, S. R., C. Becker, and C. E. Caceres. 2007. “Parasitic Castration: A Perspective from a Model of Dynamic Energy Budgets.” Integrative and Comparative Biology 47 (2): 295–309. 10.1093/icb/icm057.

Hall, Spencer R., Claes R. Becker, Meghan A. Duffy, and Carla E. Cáceres. 2012. “A Power– Efficiency Trade□off in Resource Use Alters Epidemiological Relationships.” Ecology 93 (3): 645–56. 10.1890/11-0984.1.

Hall, Spencer R., Christine J. Knight, Claes R. Becker, Meghan A. Duffy, Alan J. Tessier, and Carla E. Cáceres. 2009. “Quality Matters: Resource Quality for Hosts and the Timing of Epidemics.” Ecology Letters 12 (2): 118–28. 10.1111/j.1461-0248.2008.01264.x.

Hall, Spencer R., Meghan A. Duffy, and Carla E. Cáceres. 2005. “Selective Predation and Productivity Jointly Drive Complex Behavior in Host□Parasite Systems.” The American Naturalist 165 (1): 70–81. 10.1086/426601.

Hall, Spencer R., Joseph L. Simonis, Roger M. Nisbet, Alan J. Tessier, and Carla E. Cáceres. 2009. “Resource Ecology of Virulence in a Planktonic Host□Parasite System: An Explanation Using Dynamic Energy Budgets.” The American Naturalist 174 (2): 149–62. 10.1086/600086.

Harjoe, Carmen C., Julia C. Buck, Jason R. Rohr, Claire E. Roberts, Deanna H. Olson, and Andrew R. Blaustein. 2022. “Pathogenic Fungus Causes Density□ and Trait□mediated Trophic Cascades in an Aquatic Community.” Ecosphere 13 (4). 10.1002/ecs2.4043.

Hastings, A., Hom, C. L., Ellner, S., Turchin, P., & Godfray, H. C. J. (1993). Chaos in ecology: is mother nature a strange attractor?. Annual review of ecology and systematics, 1–33.

Hilker, Frank M., and Kirsten Schmitz. 2008. “Disease-Induced Stabilization of Predator–Prey Oscillations.” Journal of Theoretical Biology 255 (3): 299–306. 10.1016/j.jtbi.2008.08.018.

Hilker, Frank M., Michel Langlais, and Horst Malchow. 2009. “The Allee Effect and Infectious Diseases: Extinction, Multistability, and the (Dis□)Appearance of Oscillations.” The American Naturalist 173 (1): 72–88. 10.1086/593357.

Hite, Jessica L., Rachel M. Penczykowski, Marta S. Shocket, et al. 2017. “Allocation, Not Male Resistance, Increases Male Frequency during Epidemics: A Case Study in Facultatively Sexual Hosts.” Ecology 98 (11): 2773–83. 10.1002/ecy.1976.

Hite, Jessica L., Alaina C. Pfenning, and Clayton E. Cressler. 2020. “Starving the Enemy? Feeding Behavior Shapes Host-Parasite Interactions.” Trends in Ecology & Evolution 35 (1): 68–80. 10.1016/j.tree.2019.08.004.

Holling, C. S. 1959. “Some Characteristics of Simple Types of Predation and Parasitism.” The Canadian Entomologist 91 (7): 385–98. 10.4039/ent91385-7.

Hudson, Peter J., Andrew P. Dobson, and Kevin D. Lafferty. 2006. “Is a Healthy Ecosystem One That Is Rich in Parasites?” Trends in Ecology & Evolution 21 (7): 381–85. 10.1016/j.tree.2006.04.007.

Hurtado, Paul J., Spencer R. Hall, and Stephen P. Ellner. 2014. “Infectious Disease in Consumer Populations: Dynamic Consequences of Resource-Mediated Transmission and Infectiousness.” Theoretical Ecology 7 (2): 163–79. 10.1007/s12080-013-0208-2.

Johnson, Pieter T.J., Andrew Dobson, Kevin D. Lafferty, et al. 2010. “When Parasites Become Prey: Ecological and Epidemiological Significance of Eating Parasites.” Trends in Ecology & Evolution 25 (6): 362–71. 10.1016/j.tree.2010.01.005.

Keesing, Felicia, and Richard S. Ostfeld. 2021. “Dilution Effects in Disease Ecology.” Ecology Letters 24 (11): 2490–505. 10.1111/ele.13875.

Klausmeier, Christopher A. 2008. “Floquet Theory: A Useful Tool for Understanding Nonequilibrium Dynamics.” Theoretical Ecology 1 (3): 153–61. 10.1007/s12080-008-0016-2.

Kretzschmar, M., R.M. Nisbet, and E. Mccauley. 1993. “A Predator-Prey Model for Zooplankton Grazing on Competing Algal Populations.” Theoretical Population Biology 44 (1): 32–66. 10.1006/tpbi.1993.1017.

Lafferty, Kevin D., Stefano Allesina, Matias Arim, et al. 2008. “Parasites in Food Webs: The Ultimate Missing Links.” Ecology Letters 11 (6): 533–46. 10.1111/j.1461-0248.2008.01174.x.

Lafferty, Kevin D., Andrew P. Dobson, and Armand M. Kuris. 2006. “Parasites Dominate Food Web Links.” Proceedings of the National Academy of Sciences 103 (30): 11211–16. 10.1073/pnas.0604755103.

Lafferty, Kevin D., and Armand M. Kuris. 2009. “Parasitic Castration: The Evolution and Ecology of Body Snatchers.” Trends in Parasitology 25 (12): 564–72. 10.1016/j.pt.2009.09.003.

Lever, J. J., Van Nes, E. H., Scheffer, M., & Bascompte, J. (2023). Five fundamental ways in which complex food webs may spiral out of control. Ecology Letters, 26(10), 1765–1779. 10.1111/ele.14293

Levins, Richard. 1974. “DISCUSSION PAPER: THE QUALITATIVE ANALYSIS OF PARTIALLY SPECIFIED SYSTEMS.” Annals of the New York Academy of Sciences 231 (1): 123–38. 10.1111/j.1749-6632.1974.tb20562.x.

McCann, Kevin, Alan Hastings, and Gary R. Huxel. 1998. “Weak Trophic Interactions and the Balance of Nature.” Nature 395 (6704): 794–98. 10.1038/27427.

Murdoch, William W., Cheryl J. Briggs, and Roger M. Nisbet. 2003. Consumer-Resource Dynamics. Online-Ausg. Monographs in Population Biology 36. Princeton University Press.

Novak, Mark, Justin D. Yeakel, Andrew E. Noble, et al. 2016. “Characterizing Species Interactions to Understand Press Perturbations: What Is the Community Matrix?” Annual Review of Ecology, Evolution, and Systematics 47 (1): 409–32. 10.1146/annurev-ecolsys-032416-010215.

Penczykowski, Rachel M., Brian C. P. Lemanski, R. Drew Sieg, et al. 2014. “Poor Resource Quality Lowers Transmission Potential by Changing Foraging Behaviour.” Functional Ecology 28 (5): 1245–55. 10.1111/1365-2435.12238.

Penczykowski, Rachel M., Marta S. Shocket, Jessica Housley Ochs, et al. 2022. “Virulent Disease Epidemics Can Increase Host Density by Depressing Foraging of Hosts.” The American Naturalist 199 (1): 75–90. 10.1086/717175.

Puccia, Charles J., and Levins, Richard. 1985. “Qualitative Modeling of Complex Systems.” Harvard University Press. 10.4159/harvard.9780674435070.c6.

Ramesh, Ashwini, and Spencer R. Hall. 2023. “Niche Theory for Within□host Parasite Dynamics: Analogies to Food Web Modules via Feedback Loops.” Ecology Letters 26 (3): 351–68. 10.1111/ele.14142.

Ramesh, Ashwini, and Spencer R. Hall. 2025. In press.

Ranjit, Buddhadev, Arnab Chattopadhyay, Arindam Mandal, Santosh Biswas, and Joydev Chattopadhyay. 2025. “Beyond Predation: Fish–Coral Interactions Can Tip the Scales of Coral Disease.” Journal of Theoretical Biology 599 (February): 112031. 10.1016/j.jtbi.2024.112031.

Rosenzweig, M. L., and R. H. MacArthur. 1963. “Graphical Representation and Stability Conditions of Predator-Prey Interactions.” The American Naturalist 97 (895): 209–23. 10.1086/282272.

Rosenzweig, Michael L. 1971. “Paradox of Enrichment: Destabilization of Exploitation Ecosystems in Ecological Time.” Science 171 (3969): 385–87. 10.1126/science.171.3969.385.

Saifuddin, Md., Santanu Biswas, Sudip Samanta, Susmita Sarkar, and Joydev Chattopadhyay. 2016. “Complex Dynamics of an Eco-Epidemiological Model with Different Competition Coefficients and Weak Allee in the Predator.” Chaos, Solitons & Fractals 91 (October): 270–85. 10.1016/j.chaos.2016.06.009.

Sarnelle, Orlando, and Alan E. Wilson. 2008. “TYPE III FUNCTIONAL RESPONSE INDAPHNIA.” Ecology 89 (6): 1723–32. 10.1890/07-0935.1.

Schatz, Greg S., and Edward McCauley. 2007. “Foraging Behavior by Daphnia in Stoichiometric Gradients of Food Quality.” Oecologia 153 (4): 1021–30. 10.1007/s00442-007-0793-0.

Scheffer, Martin. 2009. “Alternative stable states and regime shifts in ecosystems.” In The Princeton Guide to Ecology: 395–406. Princeton university press.

Schröder, Arne, Lennart Persson, and André M. De Roos. 2005. “Direct Experimental Evidence for Alternative Stable States: A Review.” Oikos 110 (1): 3–19. 10.1111/j.0030-1299.2005.13962.x.

Simon, Margaret W., Michael Barfield, and Robert D. Holt. 2022. “When Growing Pains and Sick Days Collide: Infectious Disease Can Stabilize Host Population Oscillations Caused by Stage Structure.” Theoretical Ecology 15 (4): 285–309. 10.1007/s12080-022-00543-z.

Smith, Val H., Robert D. Holt, Marilyn S. Smith, Yafen Niu, and Michael Barfield. 2015. “Resources, Mortality, and Disease Ecology: Importance of Positive Feedbacks between Host Growth Rate and Pathogen Dynamics.” Israel Journal of Ecology and Evolution 61 (1): 37–49. 10.1080/15659801.2015.1035508.

Strauss, Alexander T., Anna M. Bowling, Meghan A. Duffy, Carla E. Cáceres, and Spencer R. Hall. 2018. “Linking Host Traits, Interactions with Competitors and Disease: Mechanistic Foundations for Disease Dilution.” Functional Ecology 32 (5): 1271–79. 10.1111/1365-2435.13066.

Strauss, Alexander T., Jessica L. Hite, David J. Civitello, Marta S. Shocket, Carla E. Cáceres, and Spencer R. Hall. 2019. “Genotypic Variation in Parasite Avoidance Behaviour and Other Mechanistic, Nonlinear Components of Transmission.” Proceedings of the Royal Society B: Biological Sciences 286 (1915): 20192164. 10.1098/rspb.2019.2164.

Strogatz, Steven H. 2024. Nonlinear Dynamics and Chaos: With Applications to Physics, Biology, Chemistry, and Engineering. 3rd ed. Chapman and Hall/CRC. 10.1201/9780429398490.

Uszko, Wojciech, Sebastian Diehl, Nadine Pitsch, Kathrin Lengfellner, and Thomas Müller. 2015. “When Is a Type III Functional Response Stabilizing? Theory and Practice of Predicting Plankton Dynamics under Enrichment.” Ecology 96 (12): 3243–56. 10.1890/15-0055.1.

Walsman, Jason Cosens, Alexander Thomas Strauss, and Spencer Ryan Hall. 2022. “Parasite□driven Cascades or Hydra Effects: Susceptibility and Foraging Depression Shape Parasite–Host–Resource Interactions.” Functional Ecology 36 (5): 1268–78. 10.1111/1365-2435.14030.

